# Hyper-adaptation in the Human Brain: Functional and structural changes in the foot section of the primary motor cortex in a top wheelchair racing Paralympian

**DOI:** 10.1101/2022.03.17.484824

**Authors:** Tomoyo Morita, Satoshi Hirose, Nodoka Kimura, Hiromasa Takemura, Minoru Asada, Eiichi Naito

## Abstract

The human brain has the capacity to drastically alter its somatotopic representations in response to congenital or acquired limb deficiencies and dysfunctions. The main purpose of the present study was to elucidate such extreme adaptability in the brain of an active top wheelchair racing Paralympian (participant P1) who has congenital paraplegia (dysfunction of bilateral lower limbs). Participant P1 has undergone long-term wheelchair racing training using bilateral upper limbs and has won a total of 19 medals in six consecutive summer Paralympic games as of 2021. We examined the functional and structural changes in the foot section of the primary motor cortex (M1) in participant P1 as compared to able-bodied control participants. We also examined the functional and structural changes in three other individuals (participants P2, P3, and P4) with acquired paraplegia, who also had long-term non-use period of the lower limbs and had undergone long-term training for wheelchair sports (but not top athletes at the level of participant P1). We measured brain activity in all the participants using functional magnetic resonance imaging (MRI) when bimanual wrist extension-flexion movement was performed, and the structural MRI images were collected. Compared to 37 control participants, participant P1 showed significantly greater activity in the M1 foot section during the bimanual task, and significant local GM expansion in this section. Significantly greater activity in the M1 foot section was also observed in participant P4, but not in P2 and P3, and the significant local GM expansion was observed in participant P2, but not in P3 and P4. Thus, functional or structural change was observed in an acquired paraplegic participant, but was not observed in all the paraplegic participants. The functional and structural changes typically observed in participant P1 may represent extreme adaptability of the human brain. We discuss the results in terms of a new idea of hyper-adaptation.

## 1 Introduction

Somatotopy is a fundamental functional structure for sensorimotor processing in the brain and is rich in plasticity. Previous neuroimaging studies have shown that the human brain has the capacity to drastically change its somatotopic representations in response to congenital or acquired limb deficiencies and dysfunction (Flor et al., 1995, 2006; Bruehlmeier et al., 1998; Lotze et al., 1999, 2001; Mikulis et al., 2002; Stoeckel et al., 2009; Hahamy et al., 2017; Dempsey-Jones et al., 2019; Hahamy and Makin, 2019; Nakagawa et al., 2020).

The main purpose of the present study was to elucidate such higher adaptability in the brain of an active top, wheelchair racing Paralympian (participant P1), who had congenital paraplegia (dysfunction of bilateral lower limbs). Participant P1 had received long-term wheelchair racing training using bilateral upper limbs, since she was eight years old, and had won a total of 19 medals in six consecutive summer Paralympic games as of 2021. We examined the functional and structural changes in the foot section of the primary motor cortex (M1) in participant P1, as compared to able-bodied control participants.

We conducted functional and structural magnetic resonance imaging (MRI). In the functional MRI experiment, prompted by the previous reports that somatotopic representation of the primary sensorimotor cortices in persons with congenital deficiency of a limb (e.g. upper limbs) is involved in sensory-motor processing of the other body parts (e.g. lower limbs; Hahamy et al., 2017; Dempsey-Jones et al., 2019; Hahamy and Makin, 2019; Nakagawa et al., 2020), we tested our hypothesis that the M1 foot section of participant P1 was involved in sensory-motor processing of the hand, which is rarely seen in able-bodied persons (Morita et al., 2021a). In the present study, we were particularly interested in the M1 because this is the executive locus of voluntary limb movement. In the structural MRI experiment, we examined the change in volume of the gray matter (GM) in the foot section of M1 of participant P1. It is known that long-term intensive hand/finger training, for manipulating musical instruments, causes GM expansion in the M1 hand section (Gaser and Schlaug, 2003). If the M1 foot section of participant P1 had been used as the hand section through her long-term training for wheelchair racing, using the upper limbs, we may expect the GM expansion (increase in GM volume) in this section. In addition to these investigations, we also explored the change in white matter (WM) in the brain of participant P1. If the GM of the M1 foot section of participant P1 is expanding, we may also expect the development of the nerve fibers that connect between the M1 foot section and other regions of the brain, resulting in expansion of WM which contains such developed nerve fibers.

The secondary purpose of the present study was to test whether the functional and structural changes expected in participant P1 are specific to this participant or common to other paraplegic participants. We tried to recruit paraplegic participants who had a long-term non-use period of the lower limbs and long-term wheelchair sports training and obtained three participants. One participant had paraplegia at the age of one (P2), and the remaining two had paraplegia due to spinal cord injury at the ages of 17 (P3) and of 21 (P4), respectively. They were acquired paraplegic persons having leg non-use period of more than 30 years and long-term training for wheelchair sports (Table 1). But none of them were top athletes at the level of participant P1.

**Table 1.**
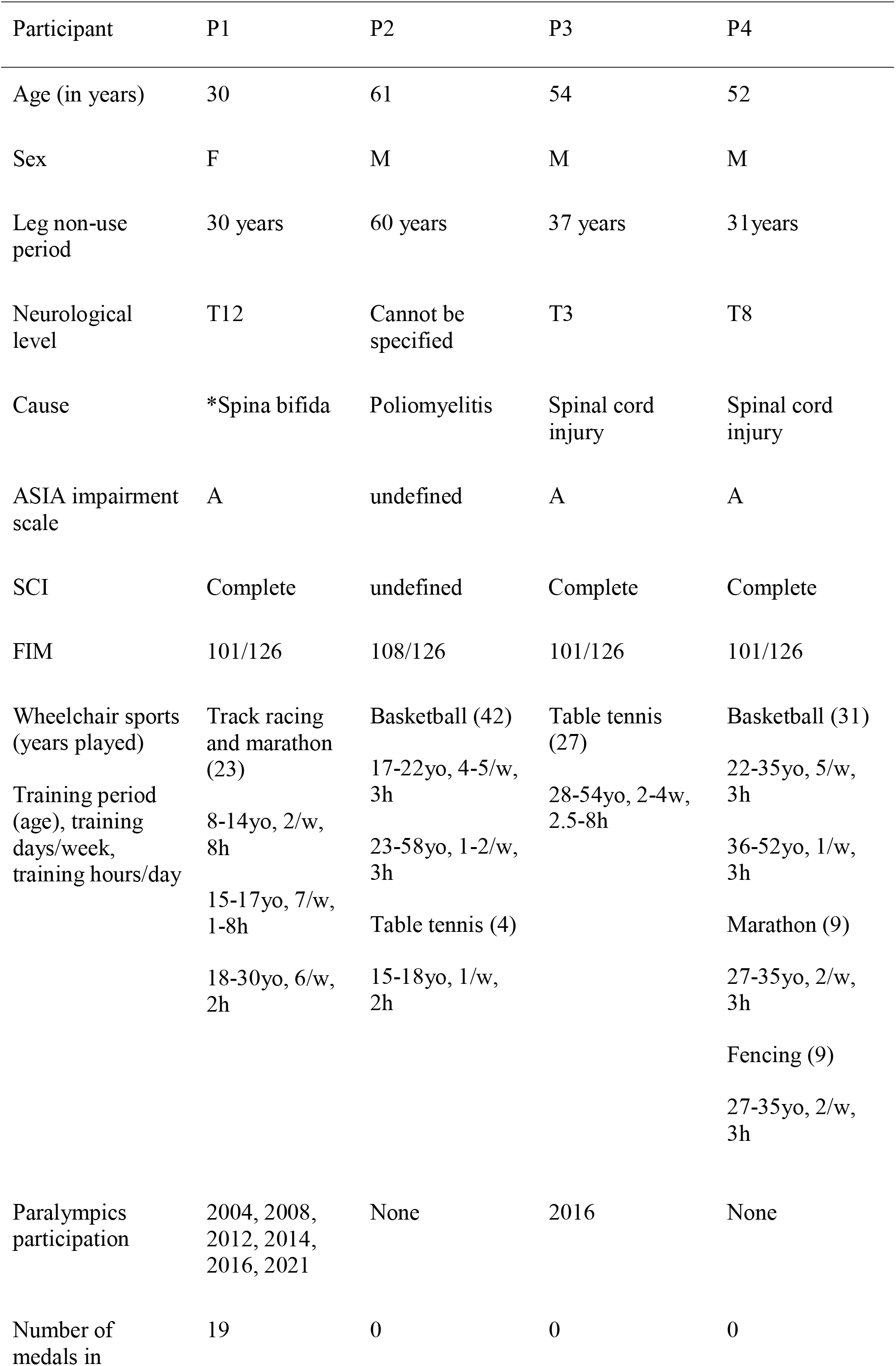
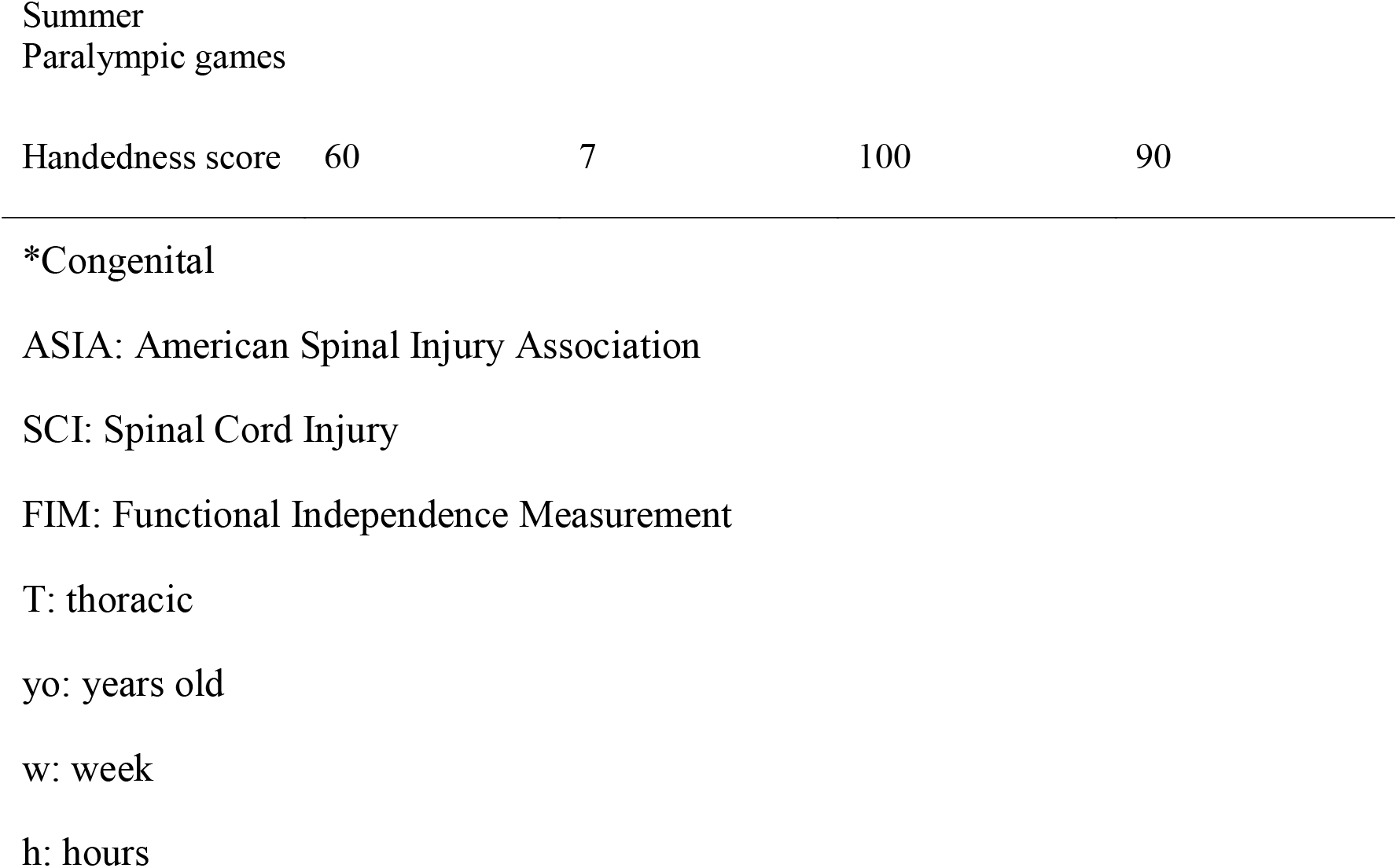
Participants’ information

In both functional and structural MRI experiments, our region of interest (ROI) was the M1 foot section. It is known that the foot section of M1 is represented in the medial wall motor region, which is closely located in the hand region of the cingulate motor area (CMA) in monkeys (He et al., 1995) and humans (Ehrsson et al., 2003; Naito et al., 2007; Amiez and Petrides, 2014). In addition, the trunk section of M1 is known to be closely located to the M1 foot section (Naito et al., 2021). Therefore, we first defined the ROI in the foot section of M1 (M1 foot ROI) using the functional MRI data obtained when 37 able-bodied control participants performed a right foot, right hand, and trunk tasks.

In the functional MRI experiment, we scanned the brain activity when the four paraplegic participants (P1, P2, P3, and P4), and the control participants performed bimanual wrist extension-flexion movements. We selected a bimanual task because moving both hands and arms simultaneously is essential movement for pedaling a wheelchair. We first directly compared the brain activity obtained from each paraplegic participant (P1, P2, P3, or P4) to that of the control participants to explore regions, in which a paraplegic participant shows significantly greater activity than the control participants, within the M1 foot ROI (contrast analysis; see below). Next, we tested if the activity obtained from the ROI and the number of activated voxels identified in the ROI were significantly greater in each paraplegic participant when compared with the control participants (ROI analysis; see below).

In the structural MRI experiment, we collected the structural MRI images from all the participants and performed voxel-based morphometry (VBM) analysis. We first explored regions, in which a paraplegic participant shows significant change in GM volume as compared to the control participants, within the M1 foot ROI (contrast analysis; see below). Next, we tested if the GM volume of the ROI in each paraplegic participant (P1, P2, P3, or P4) was significantly different from that of the control participants (ROI analysis; see below). Finally, we explored whether there was a significant change in WM volume in the whole brain space in each paraplegic participant as compared to the control participants.

## 2 Materials and Methods

### 2.1 Participants

One participant with congenital paraplegic (participant P1) and three participants with acquired paraplegia (P2, P3, and P4) participated in this study. The details of the participants are summarized in Table 1. Participant P1 was an active, top, wheelchair racing Paralympian aged 30. She began racing at the age of eight and had long-term wheelchair racing training using the bilateral upper limbs since then. She has won a total of 19 medals in the six consecutive Summer Paralympic Games as of 2021. Participants P2, P3, and P4 were not top athletes at the level of participant P1, but they had more than 30 years of leg non-use period and long-term wheelchair sports training (Table 1). Participant P2 had paraplegia at the age of one and had 42 years of experience in wheelchair basketball and 4 years of experience in wheelchair table tennis. Participant P3 had paraplegia at the age of 17 and had 27 years of experience in wheelchair table tennis. He was a Paralympian in Río de Janeiro. Participant P4 had paraplegia at the age of 21, and had 31 years of experience in wheelchair basketball, 9 years of experience in wheelchair marathon, and 9 years of experience in wheelchair fencing. Participants P1, P3, and P4 had no somatic sensations (light touch and pin prick) from their lower limbs and had complete immobility. Participant P2 had complete immobility of his lower limbs; nonetheless, there were somatic sensations (light touch and pin prick). These were evaluated by a physiotherapist with more than 10 years of experience (NK, one of the authors). We confirmed the handedness of the participants using the Edinburgh Handedness Inventory (Oldfield, 1971), and participants P1, P3, and P4 were right-handers, and participant P2 was ambidextrous (Table 1).

Regarding the control participants, we recruited right-handed and -footed able-bodied adults (n = 37; 37.4 ± 10.9 [mean ± standard deviation] years old, range 25-59 years old, 25 female). They had experience in various sports since their school days, but none of them were athletes participating in a particular sport. We confirmed the handedness of the control participants using the Edinburgh Handedness Inventory (95.9 ± 7.6, Oldfield, 1971). Moreover, we determined the dominant operating foot using a question “which leg would you use to kick a ball” in the Inventory. This question is shown to be valid to determine the dominant operating foot (Van Melick et al. 2017). Among the participants, 33 participants reported using their right leg and 4 used both legs. Based on self-reports, none of the participants had a history of neurological, psychiatric, or movement disorders.

The study protocol was approved by the Ethics Committee of the National Institute of Information and Communications Technology (NICT) and by the MRI Safety Committee of the Center for Information and Neural Networks (CiNet; no. 2003260010). We explained the details of the experiment to each participant before the experiment, after which they provided written informed consent. The study was conducted according to the principles and guidelines of the Declaration of Helsinki (1975).

### 2.2 General Procedure

We conducted both functional and structural MRI experiments. We first collected the functional data, and at the end of entire experiment, we collected the structural MRI data from all participants. In the functional MRI experiment, to define the foot section of M1, the 37 control participants performed right foot, right hand, and trunk tasks (Figure 1(A)), in addition to a main bimanual task (Figure 2(A)). The paraplegic participants performed the bimanual task. After the bimanual task, both the control and paraplegic participants also performed right-hand active and passive movement tasks, which we intend to report in a future paper.

**Figure 1.**
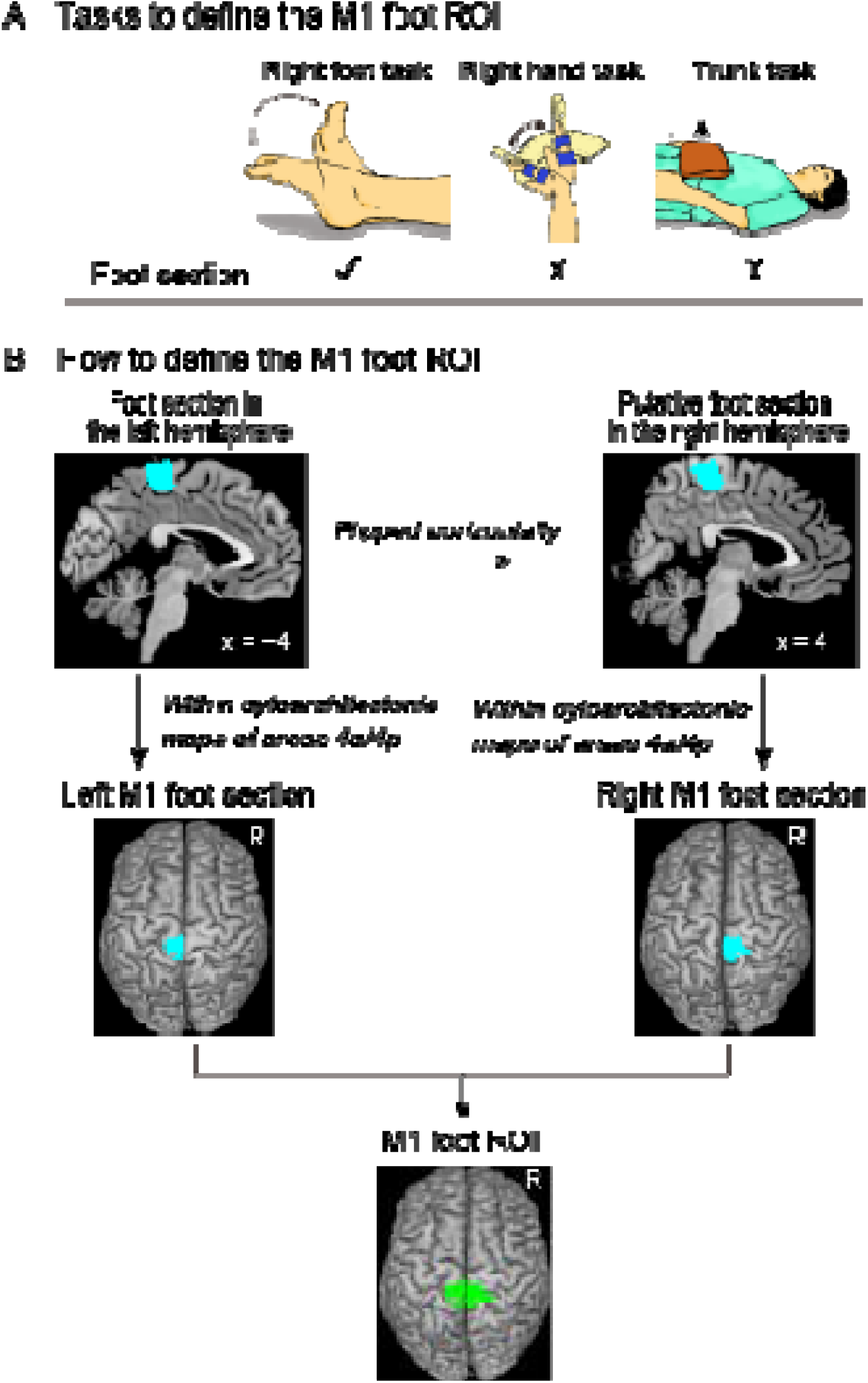
Defining the M1 foot ROI. (A): Tasks to define the M1 foot ROI. We use the right foot, right hand, and trunk tasks to identify the brain regions that are activated during the right foot task, but not during the right hand and trunk tasks. (B): Procedures to define the M1 foot ROI (green section). Please see the text for details. Abbreviations: M1, primary motor cortex; ROI, region-of-interest.

**Figure 2.**
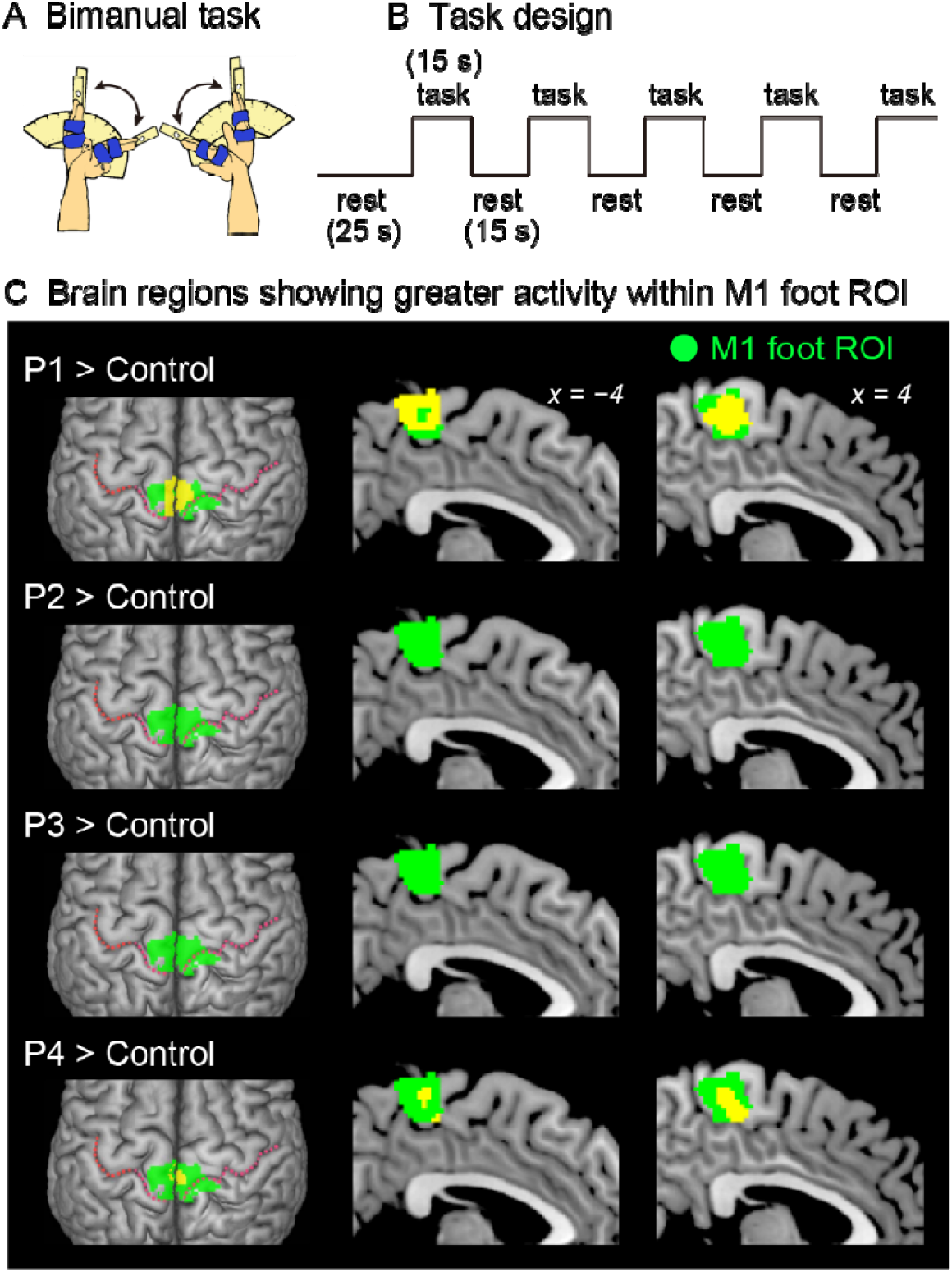
Results from contrast analysis for functional data. (A): Bimanual task used in the present study. All of the participants continuously exert cyclic in-phase extension–flexion movements of their left and right wrists in synchronization with 1-Hz tones. (B): Task design of the bimanual task. Please see the text for details. (C): Results from one-to-many two-sample t-test. Significant clusters of voxels (yellow sections) showing greater activity within the M1 foot ROI when compared with the control participants are shown. Participants P1 and P4 showed a significant cluster (yellow sections) in the M1 foot ROI (green sections). Participant P2 and P3 had no significant cluster. These sections are superimposed on the MNI standard brain. The top view, and sagittal slices (x = −4 and 4) are shown in each row. In each top view panel, pink dotted lines indicate the central sulci. Abbreviations: M1, primary motor cortex; ROI, region-of-interest; MNI, Montreal Neurological Institute.

Before the fMRI experiment, we explained the tasks to be performed in the scanner to every participant, and they experienced the tasks outside the MRI room to familiarize themselves with the tasks. Thereafter, the participants entered the room and were placed in the MRI scanner. Their heads were immobilized using sponge cushions and an adhesive tape, and their ears were plugged. The participant’s body parts (chest, pelvis, and shin) of the participants were fixed to the MRI bed using Velcro to reduce their body movements during the task. When performing a task, the participants were asked to close their eyes, relax their entire body, refrain from producing unnecessary movements, and only think of the assigned task.

Each participant completed one experimental 160-s run for each task. The run comprised five task epochs, each lasting 15 s (Figure 2(B)). Considering each epoch, the participants continuously exerted cyclic movements for each task in synchronization with cyclic audio tones. The details of each task are described below. The task epochs were separated by 15-s baseline (rest) periods. Each run also included a 25-s baseline period before the start of the first epoch. During the experimental run, we provided the participants with auditory instructions that indicated the start of a task epoch (three, two, one, start). We also provided a ‘stop’ instruction generated by a computer to notify the participants of the end of each epoch. The participants heard the same cyclic audio tones; however, they did not generate any movement during the rest periods. All the auditory stimuli were provided through an MRI-compatible headphone. An experimenter who stood beside the scanner bed checked if the participants were performing each task properly by visual inspection throughout the run.

### 2.3 MRI Data Acquisition

Functional MRI images were acquired using T2*-weighted gradient echo-planar imaging (EPI) sequences with a 3.0-Tesla MRI scanner (MAGNETOM Trio Tim; Siemens, Germany) and 32-channel array coil. We used a multiband imaging technique (multiband factor = 3), which was used in our previous study (Amemiya et al., 2021). Each volume consisted of 48 slices (slice thickness = 3.0 mm) acquired in an interleaved manner, covering the entire brain. The time interval between the successive acquisitions from the same slice was 1000 ms. An echo time of 27 ms and a flip angle of 60 ° were used. The field of view was 192 × 192 mm, and the matrix size was 64 × 64 pixels. The voxel dimensions were 3 × 3 × 3 mm along the x-, y-, and z-axes, respectively. We collected 160 volumes for each experimental run.

Regarding the structural MRI image, a T1-weighted magnetization-prepared rapid gradient echo (MP-RAGE) image was acquired using the same scanner for each participant, which was used in the following voxel-based morphometry (VBM) analysis. The imaging parameters were as follows: TR =1900 ms, TE = 2.48 ms; FA = 9 °, field of view = 256 × 256 mm^2^, matrix size =256 × 256 pixels, slice thickness = 1.0 mm; voxel size = 1 × 1 × 1 mm^3^, and 208 contiguous transverse slices.

### 2.4 Tasks to Functionally Define the Foot Section of M1

We prepared the following three tasks (Figure 1(A)) to functionally define the foot section of M1 in the control participants.

#### 2.4.1 Right foot task

The control participants performed alternating dorsi- and plantar-flexions of the right foot at a frequency of 1 Hz (left panel in Figure 1(A)). They were required to continuously perform these movements in synchronization with 1-Hz cyclic tones while relaxing their left foot. A supporter was placed under the right calf and the right leg was lifted off the bed to enable the participants to generate these movements without their right heel touching the bed. The participants were instructed to perform right foot movements within the range of their maximum dorsi- and plantar-flexion angles. These were measured outside the scanner (average range of motion across participants was approximately 68.6 ± 15.5 °). This task was used to identify the brain regions that were associated with foot movement.

#### 2.4.2 Right hand task

The control participants continuously exerted cyclic extension–flexion movements of their right wrist in synchronization with 1-Hz cyclic tones. We prepared a device to control the range of wrist motion (middle panel in Figure 1(A)), which was used in our previous study (Morita et al., 2021a). A movable hand-rest was mounted on the device (middle panel in Figure 1(A)), and the hand was fixed on the hand-rest that indicated the wrist angle. Two stoppers were fixed onto the device to control the range of the wrist motion across the task epochs and participants. They were positioned to prevent the wrist from extending beyond the straight (0 °) position and flexing beyond 60 °. The participants had to touch one of the stoppers (0 ° or 60 °) alternately with the hand-rest in synchronization with the 1-Hz audio tones while making controlled and continuous wrist extension–flexion movements. An example of kinematic data when performing this task is shown in (Morita et al., 2021a). This task was used to depict foot-specific M1 section distinctly from the CMA hand region. This is because the foot section of M1 and the CMA hand region are closely represented in the medial wall (see Introduction).

#### 2.4.3 Trunk task

The control participants pushed up a 4-kg weight placed on their abdomen (right panel in Figure 1(A)). They were asked to repeatedly perform a set of push-ups and immediate relaxation in synchronization with 0.8 Hz of tones, without moving their heads, and 0.8 Hz was chosen because in our pilot experiment, some participants reported that 1-Hz was too fast. They indicated that 0.8 Hz was comfortable to follow the movements. The participants were instructed to keep their breathing as normal as possible during the scanning. The weight was placed on their abdomen at the start of each task epoch started and removed when the epoch was completed. This task was used to identify the foot-specific M1 section, distinctly from the M1 trunk section, because the foot movement could potentially co-activated the M1 trunk section (Naito et al., 2021).

#### 2.4.4 Bimanual task (main task)

All the participants continuously exerted cyclic extension–flexion movements of their left and right wrists in synchronization with the 1-Hz tones (Figure 2(A)). The participants generated in-phase extension–flexion movements of both hands. The range of the wrist motion was between 0 ° and 60 ° as shown in the right-hand task (see above). To control the range of motion, the same device, used in the right hand task, was used for each of the left and right hands.

### 2.5 fMRI Data Preprocessing and Single-subject Analysis

To eliminate the effects of unsteady magnetization during the tasks, the first ten EPI images in each fMRI run were discarded. The imaging data were analyzed using SPM 12 (Wellcome Centre for Human Neuroimaging, London, UK) implemented in MATLAB (MathWorks, Sherborn, MA, USA). The following preprocessing was done for each participant. SPM default parameters were used unless otherwise specified. First, all of the EPI images were aligned to the first EPI image of the first session with six degrees-of-freedom (translation and rotation about x-, y-, z-axes) rigid displacement. Through this realignment procedure, we obtained the data related to the position of the head that changed over time from the first frame and through the six parameters. All the participants had a maximum displacement of less than 1.5 mm in the x-, y-, or z-plane and less than 0.1° of angular rotation about each axis during each fMRI run. These values were comparable to those from our previous study (see Morita et al., 2019), and thus no data were excluded from the analysis. The T1-weighted structural image of each participant was co-registered to the mean image of all the realigned EPI images by using affine transformation. Finally, the structural image and the realigned EPI images were spatially normalized to the standard stereotactic Montreal Neurological Institute (MNI) space (Evans et al., 1994). Normalization parameters to align the structural image to MNI template brain were calculated using the SPM12 normalization algorithm. The same parameters were used to transform the realigned EPI images. The normalized EPI images were resliced to 2-mm isotropic resolution, and the successful alignment was visually checked. Finally, the normalized images were spatially smoothed using a Gaussian kernel with a full width at half maximum of 4 mm along the x-, y-, and z-axes.

Following the preprocessing, we used a general linear model (Friston et al., 1995; Worsley and Friston, 1995) to analyze the fMRI data. We prepared a design matrix for each participant. Considering this single-subject analysis, the design matrix contained a boxcar function for the task epoch in the run, which was convolved with a canonical hemodynamic response function. To correct the residual motion-related variance after the realignment, the six realignment parameters were also included in the design matrix as regressors of no interest. In the analysis, we did not perform global mean scaling to avoid inducing Type I errors in the evaluation of negative blood oxygenation level-dependent (BOLD) responses (deactivation) (Aguirre et al., 1998). We generated an image showing the task-related activity in each task for each participant, which was used in the subsequent analyses. In this image, the effect of the cyclic tones was most likely eliminated because the participants heard the sound consistently during the task epochs and rest periods.

#### 2.5.1 Defining the M1 foot ROI

The procedure is summarized in Figure 1(B). We performed a second-level group analysis (Holmes and Friston, 1998) using a one-sample t-test to identify the brain regions that were significantly activated during the right foot task in the control participants as a whole. Here, we depicted the regions exclusively activated during the right foot task. For this purpose, we identified the brain regions that were activated during the right foot task, but not during the right hand and trunk tasks for the group of the control participants. This was carried out using two exclusive masks of images showing the right hand and trunk task-related activities. As for a mask image of the right hand task-related activity, we generated a mask image by performing one-sample t-tests to depict the brain regions that were active during the right hand task using height threshold of p < 0.05 uncorrected. The same analysis was done to generate a mask image of trunk task-related activity. We used these two mask images (union of these two images) to exclude the regions in which activity increased during the right hand and/or trunk task. We used the family wise error rate (FWE)-corrected extent threshold (p < 0.05) in the entire brain for a voxel-cluster image generated at an uncorrected height threshold of p < 0.005. We found a significant cluster of active voxels in the left (contralateral) medial wall motor regions (foot section in the left hemisphere).

To define the M1 in the medial wall motor regions, we used the cytoarchitectonic maps for areas 4a and 4p implemented in the SPM Anatomy toolbox (Eickhoff et al., 2005). We defined the left M1 foot section (the foot section of the left M1) by depicting the overlapped region between the functional cluster in the left medial wall and the cytoarchitectonic maps for areas 4a and 4p. We checked the individual activity in the left M1 foot section during the right foot, hand, and trunk tasks (see Supplementary Material and Figure S1). We also defined the right M1 foot section. We first flipped the cluster horizontally from the left medial wall to the right hemisphere. Thereafter, we defined the putative right M1 foot section by identifying the overlapped region between the flipped cluster in the right medial wall and the cytoarchitectonic maps for areas 4a and 4p. Finally, we defined the M1 foot ROI (845 voxels, voxel size = 2 × 2 × 2 mm) by combining the left and right M1 foot sections.

#### 2.5.2 Functional Analysis of bimanual task

##### 2.5.2.1 Contrast Analysis for Functional Data

First, we explored regions, in which a paraplegic participant showed significantly greater activity than the control participants during the bimanual task, within the M1 foot ROI. In this analysis, we directly compared the brain activity obtained from each paraplegic participant to that of the control participants (Figure 2(C)). This was a one-to-many two-sample t-test that was identical to Crawford and Howell t-test (Mühlau et al. 2009; Boucard et al. 2015), which is a modification of the regular independent two-sample t-test, allowing us to compare one sample with a group of multiple samples, where the within-group variance is estimated from the latter group by assuming the within-group variances in two groups are identical (Crawford and Howell, 1998). In this analysis, we included the age and sex of all participants as nuisance covariates (effect of no interest) because these factors could have an influence on the evaluation of present between-group difference. This approach has been widely used in a GLM analysis to evaluate the target factor by considering effects of the nuisance covariates (e.g., Morita et al., 2021a; Luo et al., 2014). We searched for significant clusters of voxels within the M1 foot ROI using a small volume correction (SVC; p < 0.05, FWE-corrected for a voxel-cluster image generated at an uncorrected height threshold of p < 0.005).

##### 2.5.2.2 ROI Analysis for Functional Data

In addition to the above contrast analysis, we performed ROI analysis. We first tested if the activity obtained from the M1 foot ROI during the bimanual task was significantly greater in a paraplegic participant compared to the control participants (Figure 3(A)). We calculated mean activity of the voxels in the entire M1 foot ROI in each participant. Next, we tested if the number of activated voxels identified in the ROI was significantly greater in a paraplegic participant compared to the control participants. We counted the number of activated voxels (height threshold of p < 0.005 (T > 2.61)) in the M1 foot ROI for each participant (Figure 3(B), (C)). In each ROI analysis, we performed the Crawford and Howell t-test with Bonferroni correction (Crawford and Howell, 1998; Crawford and Garthwaite, 2012) to compare the data obtained from each paraplegic participant with that of the control participants (n = 37). Given that this test was repeated for four paraplegic participants, we corrected the p-values based on the number of test repetitions (n = 4). We used the threshold of p < 0.05 with Bonferroni correction. Finally, prompted by our previous report that the M1 foot section in able-bodied persons are deactivated during hand movement (Morita et al., 2021a), we also checked for any significant decrease in activity (less than 0) in the entire M1 foot ROI during the bimanual task in the control participants as a whole by performing one-sample t-test (Figure 3(A)).

**Figure 3.**
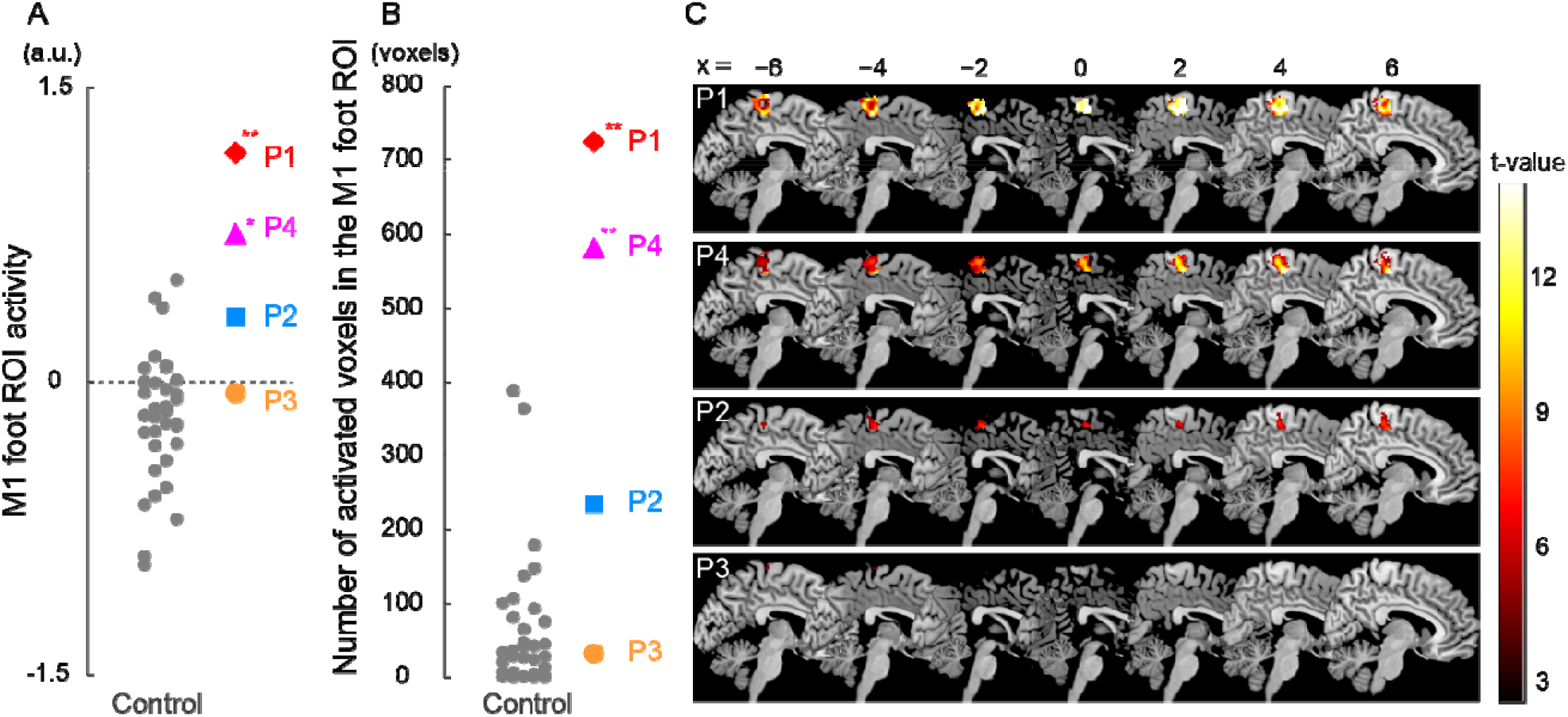
Results from ROI analysis for functional data. (A): Individual M1 foot ROI activity during the bimanual task. A vertical axis indicates the mean value of the brain activity (parameter estimates) obtained from the ROI. Participants P1 and P4 showed significantly greater M1 foot ROI activity compared to the control participants. (B): The number of activated voxels having T-value greater than 2.61 (which corresponded to height threshold p < 0.005) within the M1 foot ROI in each participant. A vertical axis indicates the number of activated voxels. The number of activated voxels in participants P1 and P4, but not in P2 and P3, were significantly larger than those in the control participants. (A, B): Red diamonds, blue rectangles, orange circles, and pink triangles represent the data obtained from paraplegic participants P1, P2, P3, and P4, respectively. Gray dots represent th data obtained from the control participants. The plotted points for the control participants are horizontally jittered to avoid over-plotting. Asterisks indicate significant differences from the control data (** *p* < 0.001, * *p* < 0.05 with Bonferroni correction). (C): Voxels having T-value greater than 2.61 (height threshold p < 0.005) within the M1 foot ROI in each participant. Please note that participants are listed in order of greater number of activated voxels, from top to bottom. These voxels are superimposed on the sagittal slices (x = −6, −4, −2, 0, 2, 4, and 6) of the MNI standard brain. Abbreviations: M1, primary motor cortex; ROI, region-of-interest; a.u., *arbitrary unit*.

#### 2.5.3 Gray matter (GM) volume analysis

##### 2.5.3.1 Contrast Analysis for GM volume

A voxel-based morphometry (VBM) analysis was performed to examine the change in GM volume in the M1 foot ROI in each paraplegic participant, compared to the control participants. First, we visually inspected the anatomical images of all the participants and confirmed the absence of observable structural abnormalities and motion artifacts. Subsequently, the data were processed using Statistical Parametric Mapping (SPM12, Wellcome Centre for Human Neuroimaging). The following steps were performed using the default settings in SPM12 as recommended by Ashburner (2010).

First, the anatomical image obtained from each participant was segmented into GM, WM, cerebrospinal fluid (CSF), and non-brain parts. Next, using Diffeomorphic Anatomical Registration Through Exponentiated Lie Algebra (DARTEL), we generated GM and WM DARTEL templates based on the anatomical images obtained from all the participants. Thereafter, we applied an affine transformation to the GM and WM DARTEL templates to align them with their tissue probability maps in the Montreal Neurological Institute (MNI) standard space. Subsequently, a segmented GM image from each participant was warped non-linearly to the GM DARTEL template in the MNI space (spatial normalization). The warped image was modulated by Jacobian determinants of the deformation field to preserve the relative GM volume, even after spatial normalization. The modulated image of each participant was smoothed with an 8-mm FWHM Gaussian kernel and resampled to a resolution of 1.5 × 1.5 × 1.5 mm^3^ voxel size. The methods described above were also used in our previous study (Morita et al., 2021b).

As in the contrast analysis for functional data, we performed one-to-many two-sample t-test by directly comparing the GM volume obtained from each paraplegic participant to that of the control participants (Figure 4(A)). This analysis has been repeatedly used in the VBM analysis (Mühlau et al., 2009; Scarpazza et al., 2013; Taubert et al., 2015). In this analysis, we included age, sex, and intracranial volume (sum of GM, WM, and CSF volumes) as nuisance covariates because these factors may have a significant effect on the results of a VBM analysis in general (Hu et al., 2011). To restrict the search volume within the GM, we used a GM mask image that was created from our present data using the SPM Masking Toolbox (Ridgway et al., 2009) (http://www0.cs.ucl.ac.uk/staff/g.ridgway/masking/). Thus, the voxels outside the mask image were excluded from the analysis. We first searched for significant clusters of voxels showing GM expansion (= increase of GM volume) within the M1 foot ROI using SVC (p < 0.05 FWE-corrected for a voxel-cluster image generated at an uncorrected height threshold of p < 0.005 (T > 2.73)). In addition, prompted by the previous reports that the GM in the primary sensory-motor cortices often shrinks after spinal cord injury even after several years of injury (Crawley et al., 2004; Freund et al., 2013; Hou et al., 2014), we also checked for significant clusters of voxels showing GM atrophy (= reduction of GM volume) within the M1 foot ROI in each paraplegic participant compared to the control participants, using the same procedure described above.

**Figure 4.**
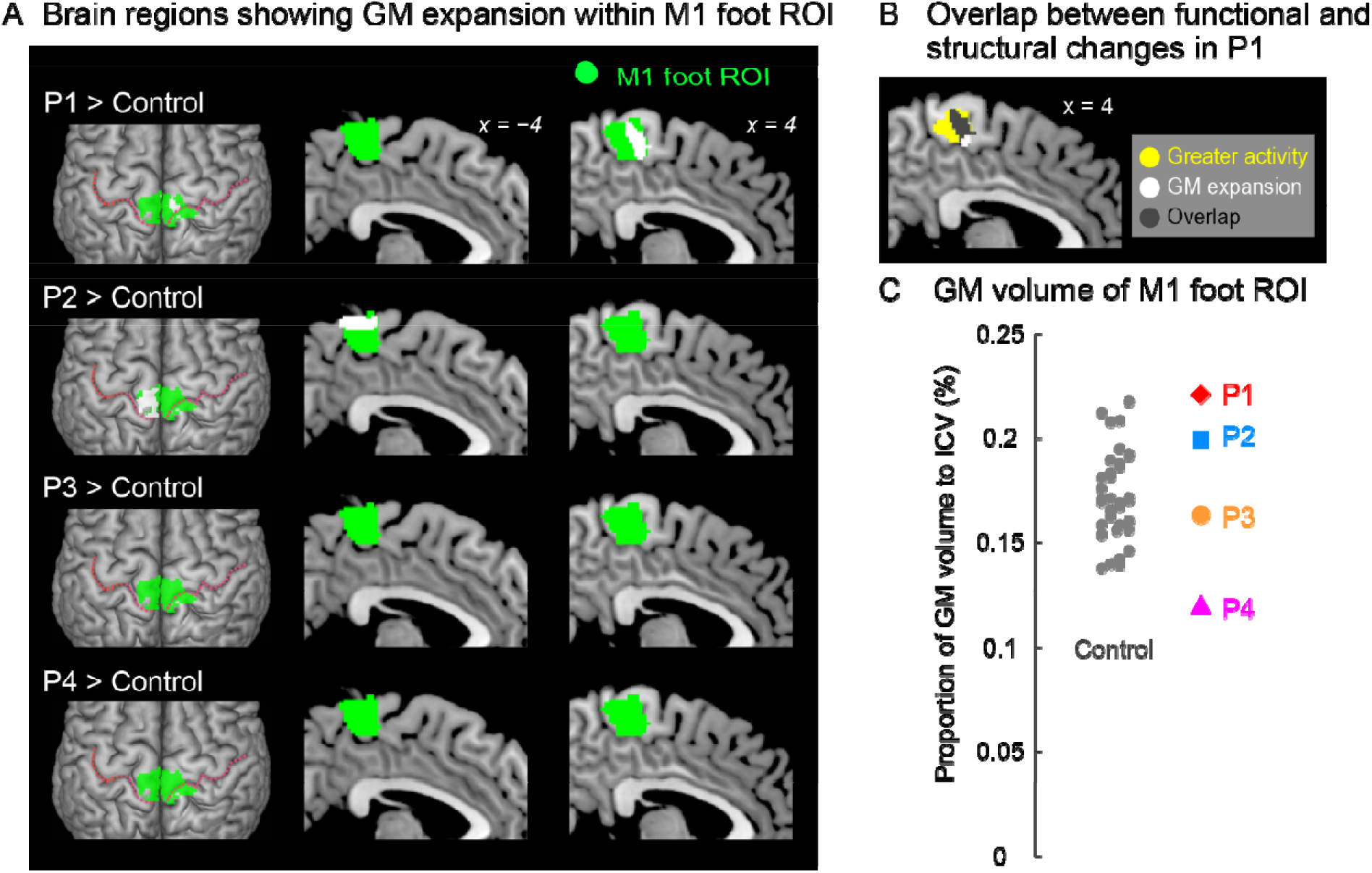
(A) Results from contrast analysis for GM volume. Significant clusters of voxels (white sections) showing GM expansion within the M1 foot ROI when compared with the control participants are shown. Only participants P1 and P2 showed a significant cluster (white sections) in the M1 foot ROI (green sections). These sections are superimposed on the MNI standard brain. The top view, and sagittal slices (x = −4 and 4) are shown in each row. In each top view panel, pink dotted lines indicate the central sulci. (B) Overlap (gray section) between the M1 foot section in which participant P1 showed significant GM expansion (white and gray sections) and the region where s e showed significantly greater activity during the bimanual task than the control participants (yellow and gray sections). (C): Individual GM volume in the M1 foot ROI. A vertical axis indicates the proportion of GM volume in the ROI to ICV. None of paraplegic participant showed significant difference from the control participants. A red diamond, blue rectangle, orange circle, and pink triangle represent the data obtained from paraplegic participants P1, P2, P3, and P4, respectively. Gray dots represent the data obtained from the control participants, and these are horizontally jittered to avoid over-plotting. Abbreviations: GM, gray matter; M1, primary motor cortex; ROI, region-of-interest; ICV, intracranial volume; MNI, Montreal Neurological Institute.

##### 2.5.3.2 ROI Analysis for GM volume

In addition to the contrast analysis, we performed a ROI analysis. We tested if the GM volume of the ROI in a paraplegic participant was significantly different (increase or decrease) from the control participants (Figure 4(C)). We calculated the GM volume of the M1 foot ROI in each participant. According to Whitwell et al. (2001), the GM volume of the M1 foot ROI was divided by intracranial volume to adjust for the size of the head of each participant. Using this value, we performed the Crawford and Howell t-test with Bonferroni correction (see above) to determine if there was significant increase or decrease in the GM volume of the entire M1 foot ROI in a paraplegic participant (two-tailed) as compared to the control participants (n = 37). Given that this test was repeated for four paraplegic participants, we corrected the p-values based on the number of test repetitions (n = 4). We used the threshold of p < 0.05 with Bonferroni correction.

#### 2.5.4 White matter (WM) volume analysis

The procedure for conducting the analysis prior to spatial normalization was identical to that in the GM volume analysis. For spatial normalization, a segmented WM image from each participant was warped non-linearly to the WM DARTEL template in the MNI space. The warped image was modulated by Jacobian determinants of the deformation field to preserve the relative WM volume, even after spatial normalization. As in the GM analysis, the modulated image of each participant was smoothed with an 8-mm FWHM Gaussian kernel and resampled to a resolution of 1.5 × 1.5 × 1.5 mm^3^ voxel size.

As we used in the GM volume analysis, one-to-many two-sample t-test was performed by directly comparing the WM volume obtained from each paraplegic participant to that of the control participants (Figure 5). We also included age, sex, and the intracranial volume as the nuisance covariates in the analysis (see above). To restrict the search volume within the WM, we used a WM mask image that was created based on our present data using the SPM Masking Toolbox (Ridgway et al., 2009). Therefore, the voxels outside this mask were excluded from the analysis. We searched for significant clusters of voxels showing WM expansion (increase of WM volume) in each paraplegic participant when compared with the control participants. Because there was no specific anatomical hypothesis available for this analysis, we explored the brain regions showing WM expansion in the entire brain space. We used the FWE-corrected extent threshold of *p* < 0.05 in the entire brain for a voxel-cluster image generated at the uncorrected height threshold of *p* < 0.005. We also checked for the brain region showing WM atrophy (decrease of WM volume) in the entire brain, in each paraplegic participant, as compared to the control participants using the procedure as described above.

**Figure 5.**
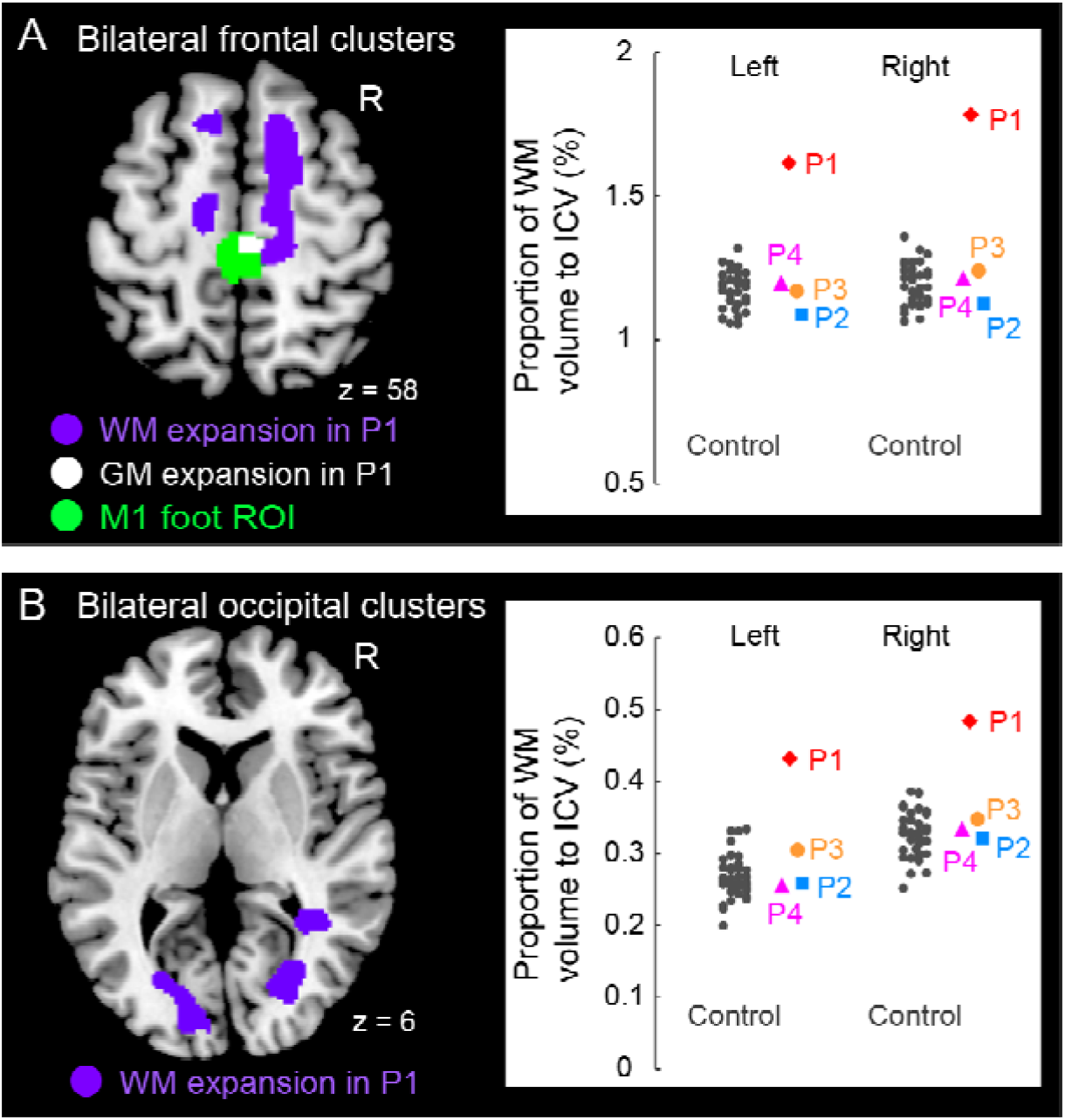
Significant white matter (WM) expansion (purple sections) in participant P1 when compared with the control participants. Significant WM expansion was observed only in participant P1. The expansions were observed from the medial prefrontal cortices to the medial-wall motor regions (A) and in the structure along with the optic radiation (B) bilaterally. These sections are superimposed on the MNI standard brain. (A): The right medial frontal cluster seemed to connect to the right M1 foot section in which she showed GM expansion (white section) within the M1 foot ROI (green section). (A, B): The right panels in A and B show scatter plots of individual WM volumes obtained from the left and right medial frontal clusters (A) and from the left and right occipital clusters (B), respectively. In each panel, a vertical axis represents the proportion of WM volume to ICV. A red diamond, blue rectangle, orange circle, and pink triangle represent the data obtained from paraplegic participants P1, P2, P3, and P4, respectively. Gray dots represent the data obtained from the control participants, and these are horizontally jittered to avoid over-plotting. Abbreviations: GM, gray matter; WM, white matter; M1, primary motor cortex; ROI, region-of-interest; ICV, intracranial volume; MNI, Montreal Neurological Institute.

To visualize the individual WM volume in the significant cluster identified in participant P1, we calculated the WM volume of the cluster in each participant. As we did in the ROI analysis for GM volume, we calculated proportion of WM volume of the significant cluster to the intracranial volume in each participant (see above). This was done purely for visualization purposes. To avoid the circular evaluation issue raised by Kriegeskorte et al. (2009), no statistical evaluations were performed.

## 3 Results

### 3.1 fMRI Results from the Bimanual Task

When the brain activity in each paraplegic participant was directly compared to that of the control participants, it was found that two of the paraplegic participants had a significant cluster of voxels showing greater activity within the M1 foot ROI (Figure 2(C)). One was participant P1, who had a significant cluster (peak coordinates: x, y, and z = 8, −24, and 62; T = 10.15, 423 voxels) in the M1 foot ROI. The other was participant P4, who also showed a significant cluster (peak coordinates: x, y, and z = 2, −32, and 64; T = 7.35; 136 voxels). Importantly, the significant clusters identified in these participants were located in the precentral region (Figure 2(C)), though the M1 foot ROI seemed to extend to the right postcentral region. Participants P2 and P3 did not show any significant clusters. Participant P2 merely had a total of 30 voxels having T-value greater than 2.73 (which corresponded to height threshold p < 0.005) within the ROI (not shown in Figure 2(C)). Participant P3 had no such voxels in the ROI.

When we looked at the individual activity obtained from the M1 foot ROI, participants P1 and P4 showed an increase in activity beyond the range of distribution of the control data (Figure 3(A)). Indeed, when we performed the Crawford and Howell t-test, participants P1 and P4 showed significantly greater activity of the M1 foot ROI as compared to the control participants (P1; t(36) = 4.06, p = 5.0 × 10 ^−4^ after Bonferroni correction, P4; t(36) = 2.82, p = 0.02 after Bonferroni correction). Neither of the participants P2 and P3 showed significantly greater M1 foot ROI activity when compared to the control participants (P2; t(36) = 1.53, p = 0.27 after Bonferroni correction, P3; t(36) = 0.33, p > 1 after Bonferroni correction). On the other hand, activity of the entire M1 foot ROI decreased (less than 0) in 26 (about 70%) of the 37 control participants (Figure 3(A)). One-sample t-test revealed significant decrease in activity in the control participants as a whole (t(36) = 3.14, p = 0.003).

Next, when the number of activated voxels identified in the M1 foot ROI (845 voxels) were counted, 724, 234, 32, and 582 voxels were found in the participant P1, P2, P3, and P4, respectively (Figure 3(B), (C)). Participants P1 and P4 showed greater number of activated voxels beyond the range of distribution of the control data (Figure 3(B)). The Crawford and Howell t-test showed that the numbers of the activated voxels in the participants P1 and P4 were significantly larger than those in the control participants (P1; t(36) = 7.46, p = 1.7 × 10^−8^ after Bonferroni correction, P4; t(36) = 5.86, p = 2.1 × 10^−6^ after Bonferroni correction). The number of the activated voxels in the participants P2 and P3 was not significantly different from those in the control participants (P2; t(36) = 1.94, p = 0.12 after Bonferroni correction, P3; t(36) = −0.33, p > 1 after Bonferroni correction). These results (the significant differences only in P1 and P4) were replicated when we counted the number of activated voxels using two different height thresholds of p < 0.01 (T > 2.35) and of p < 0.001 (T > 3.15) (not shown in Figure 3(B)).

Viewed collectively, the series of analyses consistently showed that the participants P1 and P4 had significantly greater activity in the M1 foot ROI during the bimanual task as compared to the control participants.

### 3.2 Gray matter (GM) volume change

When we explored the GM expansion (= increase in GM volume) in the M1 foot ROI in each paraplegic participant as compared to the control participants, two of the paraplegic participants (P1 and P2) showed a significant cluster of voxels showing GM expansion in the ROI (Figure 4(A)). As in the functional clusters (Figure 2(C)), the significant clusters identified in these participants were located in the precentral region. In participant P1, the significant cluster of voxels showing GM expansion was observed in the right side of the M1 foot ROI (peak coordinates = 6, −23, 54; T = 4.82; 240 voxels; Figure 4(A)). Importantly, 75% of the expanded region (gray section in Figure 4(B)) overlapped with the region in which the participant showed a significant cluster of voxels showing greater activity during the bimanual task than the control participants (yellow and gray sections in Figure 4(B)). When we calculated the dice coefficient (range from 0 to 1, with 1 meaning complete overlap) to evaluate the spatial overlap, the value was 0.29. In participant P2, the significant cluster of voxels showing GM expansion was observed in the left side of the M1 foot ROI (peak coordinates = −15, −33, 69; T = 6.10; 395 voxels; Figure 4(A)). Neither of the participants P3 and P4 showed voxels having T-value greater than 2.73 (which corresponded to height threshold p < 0.005) in the ROI.

We also examined the GM atrophy (= decrease in GM volume) in the M1 foot ROI. None of the paraplegic participants showed any significant clusters of voxels showing a decrease of GM volume within the ROI, compared to the control participants. A non-significant cluster of voxels (103 voxels having T-value greater than 2.73) was found in participant P4 (not shown in Figure 4(A)), and none of the other participants (P1, P2 and P3) had such voxels in the ROI.

When we looked at the individual GM volume of the M1 foot ROI, none of paraplegic participants showed a significant increase or decrease in GM volume when compared with the control participants, though participant P4 showed a relatively lower value of GM volume (Figure 4(C)). Thus, even though a significant cluster of voxels showing GM expansion was found in participants P1 and P2 (Figure 4(A)), the expansion was most likely localized in a limited section of the M1 foot ROI, and the GM expansion was not observed throughout the M1 foot ROI.

### 3.3 White matter (WM) volume change

Finally, when we explored the WM expansion (= increase of WM volume) in the entire brain, participant P1 showed significant clusters of voxels indicating WM expansion in the bilateral medial frontal regions (Figure 5(A)) and in the bilateral structures along the optic radiation (Figure 5(B)). Anatomical locations of these clusters are shown in Table 2. None of the other participants (P2, P3, and P4) exhibited significant WM expansion in any region of the brain. The right medial frontal cluster seemed to be connected to the right M1 foot section in which the GM expansion was observed (white section in Figure 5(A)). When we visually inspected the individual WM volume in the cluster identified in participant P1, the data obtained from participants P2, P3, and P4 were within the range of distribution of the control data, whereas participant P1 showed the largest value beyond the range of distribution of the control data in both frontal (right panel in Figure 5(A)) and occipital (right panel in Figure 5(B)) clusters of the bilateral hemispheres of the brain. Finally, none of the participants showed significant WM atrophy in any of the regions of the brain.

**Table 2.**
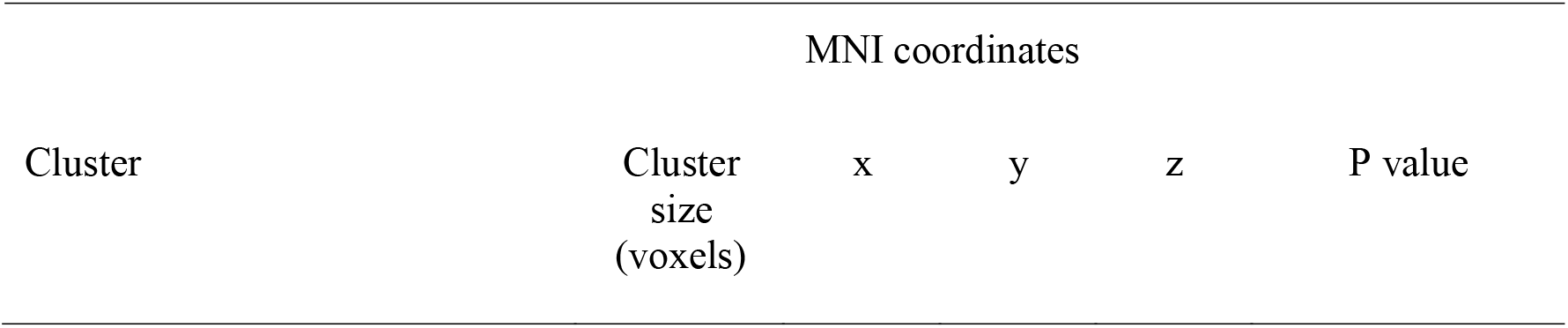
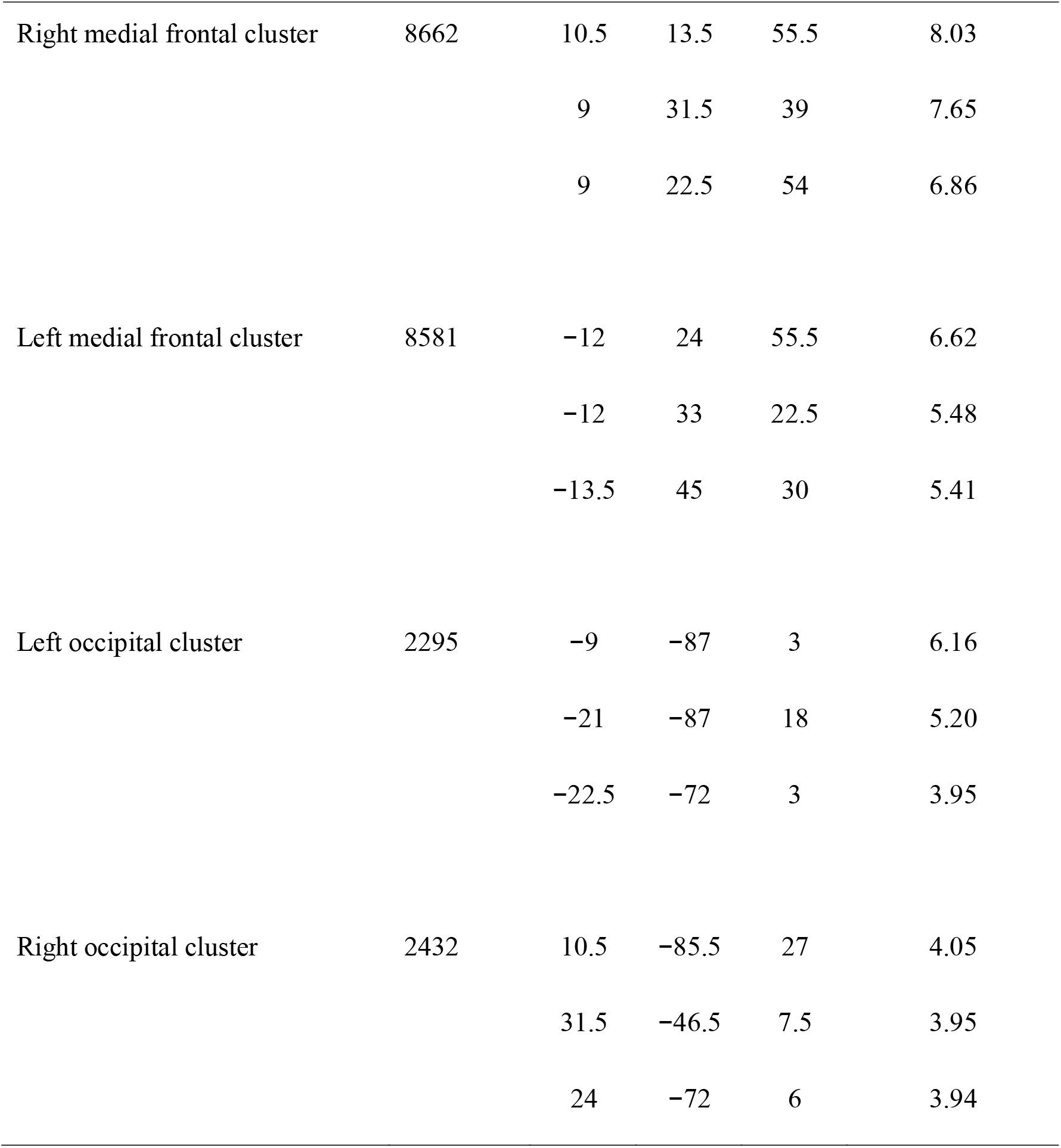
Brain regions in which participant P1 showed significant WM expansion

| Cluster | Cluster<br>size<br>(voxels) | x | y | z | P value |
| --- | --- | --- | --- | --- | --- |
| Right medial frontal cluster | 8662 | 10.5 | 13.5 | 55.5 | 8.03 |
|  |  | 9 | 31.5 | 39 | 7.65 |
|  |  | 9 | 22.5 | 54 | 6.86 |
| Left medial frontal cluster | 8581 | -12 | 24 | 55.5 | 6.62 |
|  |  | -12 | 33 | 22.5 | 5.48 |
|  |  | -13.5 | 45 | 30 | 5.41 |
| Left occipital cluster | 2295 | -9 | -87 | 3 | 6.16 |
|  |  | -21 | -87 | 18 | 5.20 |
|  |  | -22.5 | -72 | 3 | 3.95 |
| Right occipital cluster | 2432 | 10.5 | -85.5 | 27 | 4.05 |
|  |  | 31.5 | -46.5 | 7.5 | 3.95 |
|  |  | 24 | -72 | 6 | 3.94 |

## 4 Discussion

Compared to the 37 control participants, participant P1 had a significant cluster of voxels showing greater activity within the M1 foot ROI (Figure 2(C)) during the bimanual task, and a significant cluster of voxels showing GM expansion within the M1 foot ROI (Figure 4(A)). Thus, functional and structural changes were observed in participant P1 as expected. A significant cluster within the M1 foot ROI was also observed in participant P4, during the bimanual task, but not in P2 and P3 (Figure 2(C)). On the other hand, a significant cluster showing GM expansion within the ROI was observed in participant P2, but not in P3 and P4 (Figure 4(A)). Hence, the significant functional or structural change was not always observable in all the paraplegic participants, and only participant P1 exhibited both functional and structural changes in the M1 foot ROI.

There are limitations to the current study. First, we could only recruit a limited number of paraplegic participants owing to the restrictions imposed by COVID-19 although we could recruit a relatively larger number of able-bodied control participants. The number of participants was too small to conclude the difference between congenital and acquired paraplegia, the effect of the type of wheelchair sport, or the effect of the duration of wheelchair sport training. In addition, there were differences in the age and sex among the paraplegic participants. And their ages were somewhat higher than those of the control participants, though neither participant’s age was significantly different from the control group (P1; p > 1, P2; p = 0.08, P3; p = 0.29, P4; p = 0.40 after Bonferroni correction). As for the M1 foot ROI, we should have mapped the right M1 foot section by actually measuring brain activity during a left foot task. In addition, it should be borne in mind that the M1 foot section of paraplegic individuals could be different from that of the control participants in terms of its size and location. Finally, collecting more data per participant may have increased reliability of the data. Despite these limitations, this study has several implications, as discussed below.

### 4.1 Using M1 foot section for sensory-motor processing of the hand

As reported in our previous study (Morita et al., 2021a), we confirmed a significant decrease in activity of the M1 foot ROI during the bimanual task in the control participants as a whole (Figure 3(A)). Such activity decrease can be considered as cross-somatotopic inhibition (Zeharia et al., 2012; Morita et al., 2021a; Naito et al., 2021), in which the brain tries to suppress the occurrence of an unintended foot movement during hand movement. In contrast, in participant P1, we found significantly greater activity in the ROI (precentral region) during the task when compared to the control participants (Figure 2(C)). In the ROI analysis, we also confirmed significantly greater activity (Figure 3(A)) and significantly greater number of activated voxels (Figure 3(B)) in the M1 foot ROI, when compared with the control participants. These lines of evidence strongly indicate that cross-somatotopic inhibition does not exist (disinhibited) in the brain of participant P1, and that the M1 foot section is likely utilized for sensory-motor processing of the hand as if the foot section were converted to the hand section. The exact functional role of the activity in the foot section of M1 could not be determined. However, considering the fact that the hand section of M1, in persons with congenitally compromised hand functions, has a direct output to the spinal motor neurons innervating the foot muscles (Stoeckel et al., 2009), we cannot deny the possibility that the M1 foot section in participant P1 may generate motor commands to control the hand muscles.

Significantly greater activity and number of activated voxels in the M1 foot ROI were also observed in participant P4, during the bimanual task, but not in P2 and P3 (Figures 2(C) and 3). Hence, the results suggest that the use of M1 foot section for sensory-motor processing of the hand may occur even in an individual with acquired paraplegia; however, it does not always occur in all individuals with acquired paraplegia even though they have long-term non-use periods of the lower limbs and long-term wheelchair sports training.

It is known that human somatotopic representations have begun to develop from the fetal stage (Yamada et al., 2016; AboEllail et al., 2018; Dall’Orso et al., 2018), although clearly-distinct, adult-like somatotopic representations in the M1 appear to mature after 11-12 years old (Nebel et al., 2014). If we consider this evidence, we may speculate that in participant P1 with congenital paraplegia, the foot section had not developed as a foot section and developed as a section for other body parts (e.g., hand) since the fetal stage, and that in participant P2, the foot representation was still immature when he afflicted with paraplegia at the age of one.

In contrast, in the case of participant P4, the foot section would have developed as a normal foot section until being afflicted with paraplegia at the age of 21. Thus, the use of the M1 foot section for sensory-motor processing of the hand (Figures 2(C) and 3) should be considered as reorganization of the already acquired foot representation after being affected by paraplegia, and this reorganization is likely due to long-term training of upper limbs through wheelchair sports training. On the other hand, participant P3, whose foot section would also have developed as normal foot section until being afflicted with paraplegia at the age of 17, did not use the M1 foot section for sensory-motor processing of the hand (Figures 2(C) and 3). The difference between participants P3 and P4 implies the possibility that the content of wheelchair sports training (purposes of using the hands) could be an important factor to promote the use of the M1 foot section for sensory-motor processing of the hand.

From this perspective, we could point out that both participants P1 and P4 had long-term training for wheelchair track racing and marathon (Table 1) that are the sports in which the hands are used only for wheelchair mobility. The M1 foot section in an able-bodied individual is mainly used for mobility purposes (i.e., locomotion). Hence, it would be interesting to see if intensive training through the use of the hands for this purpose promotes the conversion of the M1 foot section into the hand section. However, these observations would be considered to be in the realm of speculation until they are verified by future studies.

In the case of individuals born without one hand, it has been reported that the cerebellar hand section for the missing hand, in addition to the hand section of the primary sensorimotor cortices, is involved in the sensory-motor processing of the foot (Hahamy et al., 2017; Hahamy and Makin, 2019). However, in the present study, none of our paraplegic participants showed a significant increase in the activity in the cerebellar foot section during the bimanual task (see Supplementary Material and Figure S2). This suggests the possibility of a difference in the latent potential for the conversion of somatotopic representation between the hand and foot sections in the cerebellum. In addition, it is known that long-term training of the hand (Krings et al., 2000) or foot (Naito and Hirose, 2014) movement may facilitate efficient recruitment of the brain activity (show reduction of brain activity) in its corresponding somatotopic sections. However, contrary to these previous reports, none of our paraplegic participants efficiently recruited (showed less) activity in the hand sections of the multiple motor areas (the M1, CMA, and cerebellum) during the bimanual task (see Supplementary Material and Figure S3), although they had long-term training of the upper limbs during wheelchair sports training (Table 1). Further investigations are required to generalize these findings to a larger population of individuals with long-term wheelchair sports training.

### 4.2 Gray matter (GM) changes

As expected, the contrast analysis revealed significant GM expansion within the M1 foot ROI in participant P1 when compared with the control participants (Figure 4(A)). Such significant GM expansion was also observed in participant P2, but not in P3 and P4 (Figure 4(A)). Hence, similar to the above functional changes, the GM expansion within the M1 foot ROI may occur even in an individual with acquired paraplegia, but does not always occur in all individuals with acquired paraplegia even though they have long-term non-use period of the lower limbs and long-term wheelchair sports training.

Importantly, such GM expansion was localized in a limited portion of the M1 foot ROI (Figure 4(A)), and the expansion was not observed throughout the M1 foot ROI as shown in the ROI analysis (Figure 4(C)). In addition, we could point out that the GM expansion was observed in the right precentral region in participant P1, while it was observed in the left region in participant P2, though we have no clear reasons for the difference. Although the exact physiological changes underlying GM expansion are still unknown, axon sprouting, dendritic branching synaptogenesis, neurogenesis, and changes in the glial number and morphology are suggested to be important contributors to GM expansion (Zatorre et al., 2012). Hence, our data suggest that these changes would have occurred in the localized precentral regions of participants P1 and P2.

The finding that significant GM expansion was observed in participants P1 and P2 but not in P3 and P4 (Figure 4(A)) leads us to conjecture that GM expansion is likely to occur in persons who became paraplegic at a very early stage of development. From the developmental perspective, it should also be noted that neither of the participants P3 and P4, who became paraplegic as a result of spinal cord injury during adulthood, showed significant GM atrophy in the M1 foot ROI (Figure 4(C)). Despite GM atrophy is often reported in the primary sensory-motor cortices after spinal cord injury even after several years of injury (Crawley et al., 2004; Freund et al., 2013; Hou et al., 2014). It is possible that GM atrophy would have been observed if the brains of participants P3 and P4 were scanned immediately following their spinal cord injury. Therefore, there is a possibility that long-term wheelchair sports training might have contributed to improve the putative GM atrophy. In the case of participant P4, since his M1 foot section was involved in sensory-motor processing of the hand (Figures 2(C) and 3), long-term training of the upper limbs through wheelchair sports training could have restored his putative GM atrophy. In the case of participant P3 who did not use the M1 foot section for sensory-motor processing of the hand (Figures 2(C) and 3), long-term training of other body parts through wheelchair sports training could have restored the putative atrophy.

In participant P1, 75% of the GM-expanded region (gray section in Figure 4(B)) overlapped with the region in which the participant showed a significantly greater activity than the control participants (yellow and gray sections in Figure 4(B)) during the bimanual task. Such consistency between functional and structural changes were only observed in participant P1. Hence, in participant P1, the GM was expanded in the M1 foot section that was used for sensory-motor processing of the hand. If we consider that GM expansion is deeply associated with use-dependent plasticity (Gaser and Schlaug, 2003; Draganski et al., 2004), long-term training of the upper limbs through wheelchair racing training could be an important factor for the GM expansion. We may further speculate that participant P1 has used the M1 foot section for sensory-motor processing of the hand more frequently than the control participants have used this section for foot movements. In the case of participant P2 who did not use the M1 foot section for sensory-motor processing of the hand (Figures 2(C) and 3), long-term training of other body parts through wheelchair sports training could have contributed the GM expansion (Figure 4(A)).

Finally, none of our paraplegic participants showed significant GM expansion in the hand sections of the M1, CMA, and cerebellum (see Supplementary Material and Figure S4), suggesting that our paraplegic participants use these hand sections to the same degree as the control participants use them. Similarly, none of our paraplegic participants showed significant GM expansion in the cerebellar foot section (Figure S4).

### 4.3 White matter (WM) changes

Only participant P1 showed significant WM expansion in the bilateral medial frontal regions and in the bilateral structures along the optic radiation (Figure 5). Although the exact physiological changes underlying WM expansion are not fully understood, white matter changes are thought to be related to changes in the number of axons, axon diameter, the packing density of fibers, axon branching, axon trajectories and myelination (Zatorre et al., 2012). As with GM volume, use-dependent plasticity can also be an important factor for WM expansion (Scholz et al., 2009). Thus, the long-term wheelchair racing training must be associated with the WM expansion in this participant.

WM expansion in the bilateral structures along the optic radiation are most likely associated with visual functions, which implies the possibility of the intensive use of visual system by participant P1. On the other hand, the medial frontal regions are likely to contain nerve fibers that connect the medial prefrontal cortex and the medial wall motor regions, including the M1 foot sections (probably the superior branch of the superior longitudinal fasciculus; Parlatini et al., 2017). Indeed, the right medial frontal cluster seemed to connect to the right M1 foot section in which the participant showed GM expansion (white section in Figure 5(A)). Evidence has been accumulating to support the view that the medial prefrontal cortex is important for generating highly-motivated behaviors (Walton et al., 2003; Husain and Roiser, 2018; Nakamura et al., 2021). Without a higher level of motivation, one cannot participate in six consecutive Paralympic Games and win a total of 19 medals. Thus, the WM expansion in the bilateral medial frontal regions could be associated with the repetition of highly-motivated wheelchair pedaling during long-term wheelchair racing training.

### 4.4 Hyper-adaptation

In the brain of participant P1, the M1 foot section was used for sensory-motor processing of the hand (Figures 2(C) and 3), and this claim was further corroborated by the GM expansion in the foot section that was activated during the hand sensory-motor processing (Figure 4(A), (B)). The M1 foot section is normally deactivated during the hand sensory-motor processing in able-bodied people (Figure 3(A); Morita et al., 2021). Hence, in this participant, the M1 foot section, which is normally suppressed during the hand sensory-motor processing, has drastically changed to become used for this sensory-motor processing other than its original function of foot motor control. We want to propose that such phenomenon (a brain region that is not normally involved in a certain information processing, but is rather suppressed during this information processing, in able-bodied persons, is chronically disinhibited and is regularly used during this information processing), could be better referred to hyper-adaptation. Because such adaptation may represent extreme adaptability of the human brain, which is rarely seen in typically developed individuals and should be distinct from normal adaptions as we often observe in force-field (e.g., Shadmehr and Mussa-Ivaldi et al., 1994) and visuo-motor (e.g., Imamizu et al., 2000) adaptations and so on.

The use of the foot section for the hand sensory-motor processing was also observed in participant P4 (Figures 2(C) and 3), though no supportive evidence of significant GM expansion was observed in this participant. Thus, it was likely that hyper-adaptation might also occur in this participant. In participants P2 and P3, the activity in the M1 foot section during the bimanual task was comparable to that observed in the control participants (Figures 2(C) and 3). However, this does not guarantee that hyper-adaptation does not occur in their brains because their M1 foot sections might be used for other sensory-motor processing rather than the hand, which was not directly assessed in the current work.

Based on the above definition, tactile processing in the visual cortex of some blind individuals (e.g., Sadato et al., 1996, 2002) could also be another example of hyper-adaptation because the human visual cortex is normally suppressed during sensory-motor processing in sighted people (cross-modal inhibition; Morita et al., 2019), but the visual cortex of some blind individuals has drastically changed to become involved in tactile processing other than its original function of visual processing. Hyper-adaptation may also occur when the brain manages to adapt to irreversible and severe injuries and diseases through long-term sensory, motor and cognitive trainings. Further researches will reveal what kind of adaptation should be termed hyper-adaptation, and neural mechanisms and psycho-behavioral factors that trigger and promote hyper-adaptation.

## 6 Author Contributions

Conceptualization, T.M. and E.N.; Methodology, T.M., S.H., H.T., and E.N.; Validation, T.M., S.H., N.K., H.T., M.A., and E.N.; Formal Analysis, T.M., and E.N.; Investigation, T.M., S.H., N.K., H.T., and E.N.; Writing – Original Draft Preparation, T.M., and E.N.; Writing – Review & Editing, T.M., S.H., N.K., H.T., M.A., and E.N.; Visualization, T.M.; Supervision, M.A.; Project Administration, E.N.; Funding Acquisition, T.M., and E.N.

## 8 Acknowledgments

The authors are grateful to Ms. Keiko Ueyama and CiNet MRI staff for their support during the study. We also thank the NHK Miracle Body staff, especially Ms. Yoriko Koizumi and Ms. Shiho Sakamoto for giving us the opportunity to investigate the brain of the top wheelchair racing Paralympian. We would like to thank Editage for the English language editing.

## 9 Data Availability Statement

The data that support the findings of this study are available on request from the corresponding author. The data are not publicly available because they contain information that can compromise the privacy of the participants in this study.

## 10 Institutional Review Board Statement

The study was conducted in accordance with the guidelines of the Declaration of Helsinki (1975) and was approved by the Ethics Committee of the National Institute of Information and Communications Technology and by the MRI Safety Committee of the Center for Information and Neural Networks (no 2003260010).

## 11 Informed Consent Statement

Informed consent was obtained from all participants involved in the study.

