## Supplementary materials for "Hyper-adaptation in the Human Brain: Functional and structural changes in the foot section of the primary motor cortex in a top wheelchair racing Paralympian"

*Supplementary Material*

### 1 Individual activity in the left M1 foot section during the right foot, hand, and trunk tasks

In each task, we calculated the mean activity (parameter estimates) of the voxels in the left M1 foot section (Figure 1(B)) in each participant. We plotted the data for each task. All of the control participants showed increase in activity during the right foot task, while 33 (89 %) and 19 (51 %) control participants showed activity decrease during the right hand and trunk tasks, respectively (Figure S1(A)). Thus, all of the control participants activated the left M1 foot section during the right foot task, but this section was deactivated in the majority of the participants during the right hand task, and in the half of the participants during the trunk task. Next, we counted the number of control participants who had activity (height threshold $p < 0.005$ ($T > 2.61$)) during the right foot task in each voxel of the left M1 foot section. Figure S1(B) shows the number of such control participants, as a percentage of all (37) control participants, for each voxel. More than 50% of the control participants showed such activity in most of the voxels, in the left M1foot section, during the right foot task.

### 2 Cerebellar foot ROI analysis

In the present study, in addition to the M1 foot ROI, we prepared a cerebellar foot ROI to test the possibility that the foot section of the cerebellum was involved in the sensory-motor processing of the hand in each paraplegic participant. We tested this possibility because it had been shown that the hand section of the cerebellum in congenital one-handers was involved in sensory-motor processing of the foot (Hahamy et al., 2017; Hahamy and Makin, 2019).

In the control participants as a whole, we identified cerebellar regions that were activated during the right foot task but not during the right hand and trunk tasks. Exclusive masks of images showing right hand and trunk task-related activities (height threshold $p < 0.05$, uncorrected) were used for this process. We found a cluster of active voxels (86 voxels at a height threshold of $p < 0.005$) in the right anterior cerebellar region (peak coordinates: x, y, and z = 14, -38, and -24, respectively; Lobules II, III, and IV) that were exclusively activated during the right foot task. This corresponded well with the previously reported right foot region (Grodd et al., 2001; Naito et al., 2007). We defined this region as the right cerebellar foot section. We also putatively defined the left cerebellar foot section by simply flipping the right cerebellar cluster horizontally to the left hemisphere. We define the cerebellar foot ROI (Figure S2) by combining the left and right cerebellar foot sections.

We performed contrast analysis (one-to-many two-sample t-test) to examine whether a paraplegic participant had a significant cluster of voxels showing greater activity during the bimanual task than the control participants within the cerebellar foot ROI. None of the paraplegic participants showed significant clusters within the ROI using small volume correction (SVC; $p < 0.05$ FWE-corrected for a voxel-cluster image generated at an uncorrected height threshold of $p < 0.005$). ROI analysis was also conducted to test if each paraplegic participant showed significantly greater activity of this entire ROI than the control participants (with Crawford and Howell t-test). We confirmed that, unlike the M1 foot ROI (Figures 2 and 3), none of paraplegic participants showed any significantly greater activity in the cerebellar foot ROI (Figure S2). This suggests that it is unlikely that the cerebellar foot section was involved in hand sensory-motor processing in the paraplegic participants in the present study.

### 3 Hand ROI analysis

In the present study, we also prepared hand ROIs in the M1, CMA, and cerebellum. Long-term training of the hand (Krings et al., 2000) or foot (Naito and Hirose, 2014) movement may facilitate the efficient recruitment of the activity (reduction of the activity) in its corresponding somatotopic sections.

Therefore, we set hand ROIs to test whether our paraplegic participants showed less activity in the hand sections during the bimanual task, compared to the control participants because they had long-term training of wheelchair pedaling.

Considering the M1 hand section, we identified the precentral region that was exclusively activated during the right hand task in the control participants as a whole, using exclusive masks of images showing right foot and trunk task-related activities (height threshold p < 0.05, uncorrected). We found a cluster of active voxels (1388 voxels at a height threshold of $p < 0.005$) in the left precentral gyrus. To define the M1 in this cluster, we used the cytoarchitectonic maps for areas 4a and 4p implemented in the SPM Anatomy toolbox (Eickhoff et al., 2005). We defined the left M1 hand section by depicting the overlapped region between the left precentral cluster and cytoarchitectonic maps for areas 4a and 4p. We also defined the right M1 hand section. We first flipped the left precentral cluster horizontally to the right hemisphere. Thereafter, we defined the putative right M1 hand section by identifying the overlapped region between the flipped cluster in the right precentral gyrus and cytoarchitectonic maps for areas 4a and 4p. Finally, we defined the M1 hand ROI (618 voxels; Figure S3(A)) by combining the left and right M1 hand sections.

Regarding the CMA hand section, we identified the medial wall region that was exclusively activated during the right hand task in the control participants as a whole, using the exclusive masks of images showing right foot and trunk task-related activities (height threshold p < 0.05, uncorrected). We found a cluster of active voxels (165 voxels at a height threshold of $p < 0.005$) in the left medial wall motor region. This corresponded well with the previously reported CMA hand region (Ehrsson et al., 2003; Naito et al., 2007; Amiez and Petrides, 2014). This region was defined as the left CMA hand section. We also defined the putative right CMA hand section by flipping the left medial wall cluster horizontally to the right hemisphere. We defined the CMA hand ROI (330 voxels; Figure S3(B)) by combining the left and right CMA hand sections.

Finally, as for the cerebellar hand sections, we identified the cerebellar regions that were exclusively activated during the right hand task in the control participants as a whole, using exclusive masks of images showing right foot and trunk task-related activities (height threshold p < 0.05, uncorrected). We found two clusters of active voxels in the cerebellar regions that were exclusively activated during the right-hand task. These were consistent with the previously reported right-hand regions (Grodd et al., 2001; Naito et al., 2007). One was located in the superior lobule (252 voxels at a height threshold of $p < 0.005$; peak coordinates: x, y, and z = 20, -48, and -22; Lobule VI), and the other was located in the inferior lobule (513 voxels at a height threshold of $p < 0.005$; peak coordinates: x, y, and z = 20, -56, and -52; lobule VIIIb) of the right cerebellar hemisphere. We defined these regions as the right superior- and inferior-cerebellar hand sections, respectively. We putatively defined the left cerebellar hand sections by simply flipping the above two right-sided clusters horizontally to the left hemisphere. Eventually, we defined the superior- (Figure S3(C)) and inferior-cerebellar hand ROIs (Figure S3(D)) by combining the corresponding left and right sections for each ROI.

We performed contrast analysis (one-to-many two-sample t-test) to examine whether any of the paraplegic participants had a significant cluster of voxels showing less activity during the bimanual task than the control participants within each hand ROI. None of the paraplegic participants showed any significant clusters of voxels within either ROIs (p < 0.05 FWE-corrected for a voxel-cluster image generated at an uncorrected height threshold of $p < 0.005$). ROI analysis was also conducted to test whether any of the paraplegic participants showed significantly less activity in each ROI than the control participants (with Crawford and Howell t-test). We confirmed that none of paraplegic participants showed a significantly lesser activity in any of the hand ROIs (Figure S3). Thus, although our paraplegic participants would have used their hands for a long period of time in their training for wheelchair sports (Table 1), contrary to the previous studies (see above), none of them efficiently recruit an activity in the hand sections of the multiple motor areas during the bimanual task.

**4 Gray matter (GM) volume change in other ROIs**

We also checked the change in the gray matter (GM) volume in the cerebellar foot ROI, M1 hand ROI, CMA hand ROI, superior- and inferior-cerebellar hand ROIs in each paraplegic participant, as compared to the control group. We performed contrast analysis (one-to-many two-sample t-test) to examine whether any of the paraplegic participants had a significant cluster of voxels showing an increase or decrease in the GM volume within each ROI when compared with the control participants. None of the paraplegic participants showed any significant clusters of voxels showing GM increase or decrease within either ROIs ($p < 0.05$ FWE-corrected for a voxel-cluster image generated at an uncorrected height threshold of $p < 0.005$). ROI analysis was also conducted to test whether each paraplegic participant showed a significant GM change in each ROI compared with the control participants (with Crawford and Howell t-test). None of the paraplegic participants showed any significant GM increase (expansion) or decrease (atrophy) in either ROIs (Figure S4).

### Supplementary Figures

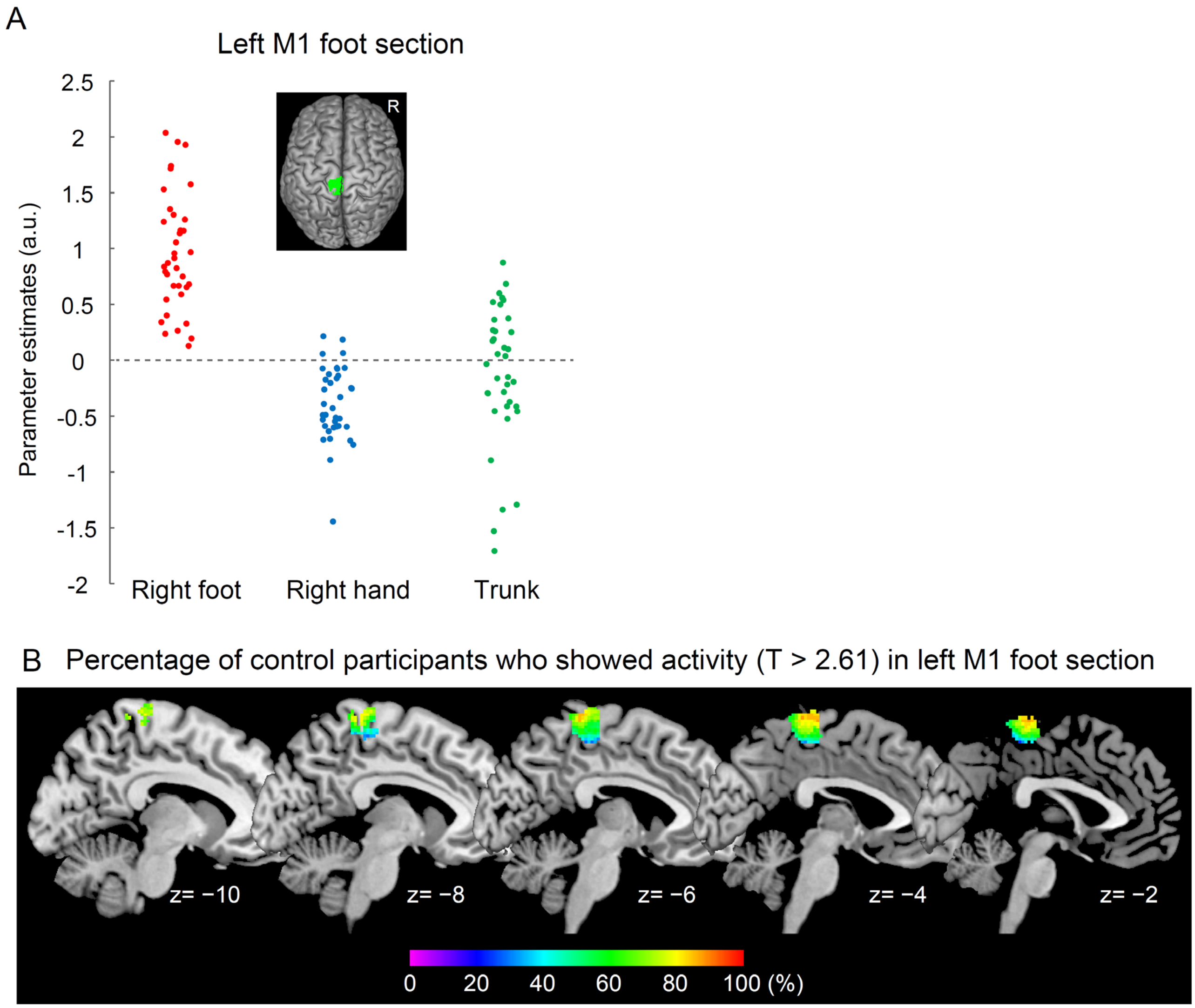


Figure S1. (A): Individual brain activity of the left M1 foot section (green section) during each of the right foot (red), right hand (blue), and trunk (green) tasks in control participants. The vertical axis indicates the mean value of brain activity (parameter estimates) of all voxels in the left foot section. (B): Color maps that indicate percentage of control participants who showed activity ($T > 2.61$) during the right foot task in the left M1 foot section. The data are superimposed on the sagittal slices ($x = -10$, $-8$, $-6$, $-4$, and $-2$) of the MNI standard brain.

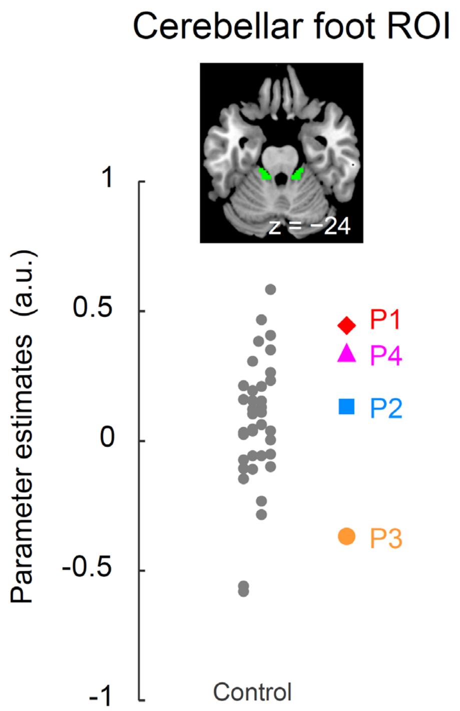


Figure S2. Individual brain activity in the cerebellar foot ROI during the bilateral task. The cerebellar foot ROI is displayed at the top. The vertical axis indicates the mean value of the brain activity (parameter estimates) of all voxels in the ROI. A red diamond, blue rectangle, orange circle, and pink triangle represent the data obtained from paraplegic participants P1, P2, P3, and P4, respectively. Gray dots represent the data obtained from the control participants. The plotted points for control participants are horizontally jittered to avoid over-plotting. All the paraplegic participants show the activity within the range of distribution of the control data. Abbreviations: ROI, region-of-interest; a.u., *arbitrary unit.*

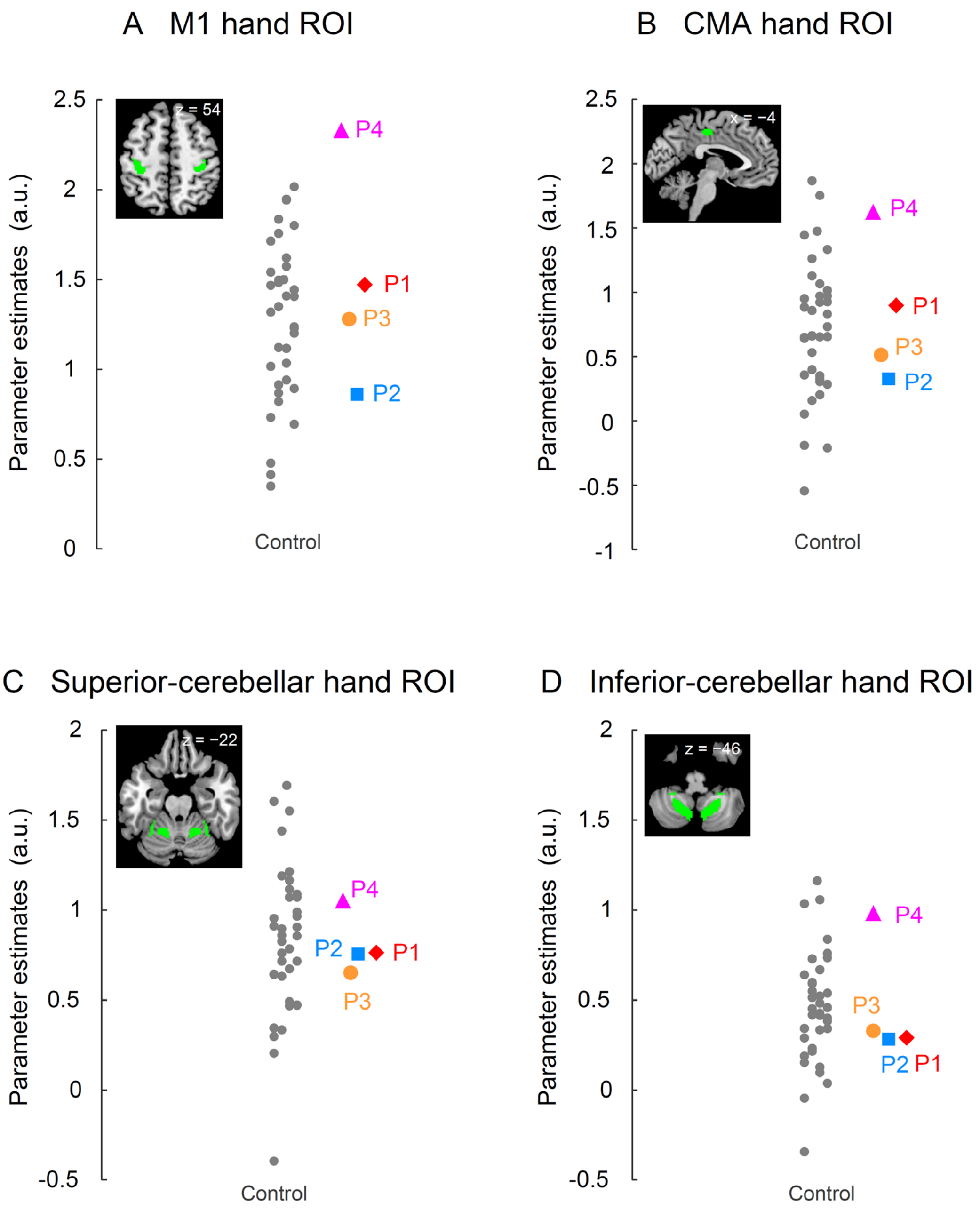


Figure S3. Individual brain activity in each of the four hand ROIs during the bilateral task. We display the data obtained from the M1 hand ROI (A), CMA hand ROI (B), superior-cerebellar hand ROI (C), and inferior-cerebellar hand ROI (D). Each ROI is displayed at the top-left of each panel. In each panel, a vertical axis indicates the mean value of the brain activity (parameter estimates) of all voxels in the ROI. Red diamonds, blue rectangles, orange circles, and pink triangles represent the data obtained from paraplegic participants P1, P2, P3, and P4, respectively. Gray dots represent the data obtained from the control participants. The plotted points are horizontally jittered to avoid over-plotting. Abbreviations: M1, primary motor cortex; CMA, cingulate motor area; ROI, region-of-interest; a.u., *arbitrary unit.*

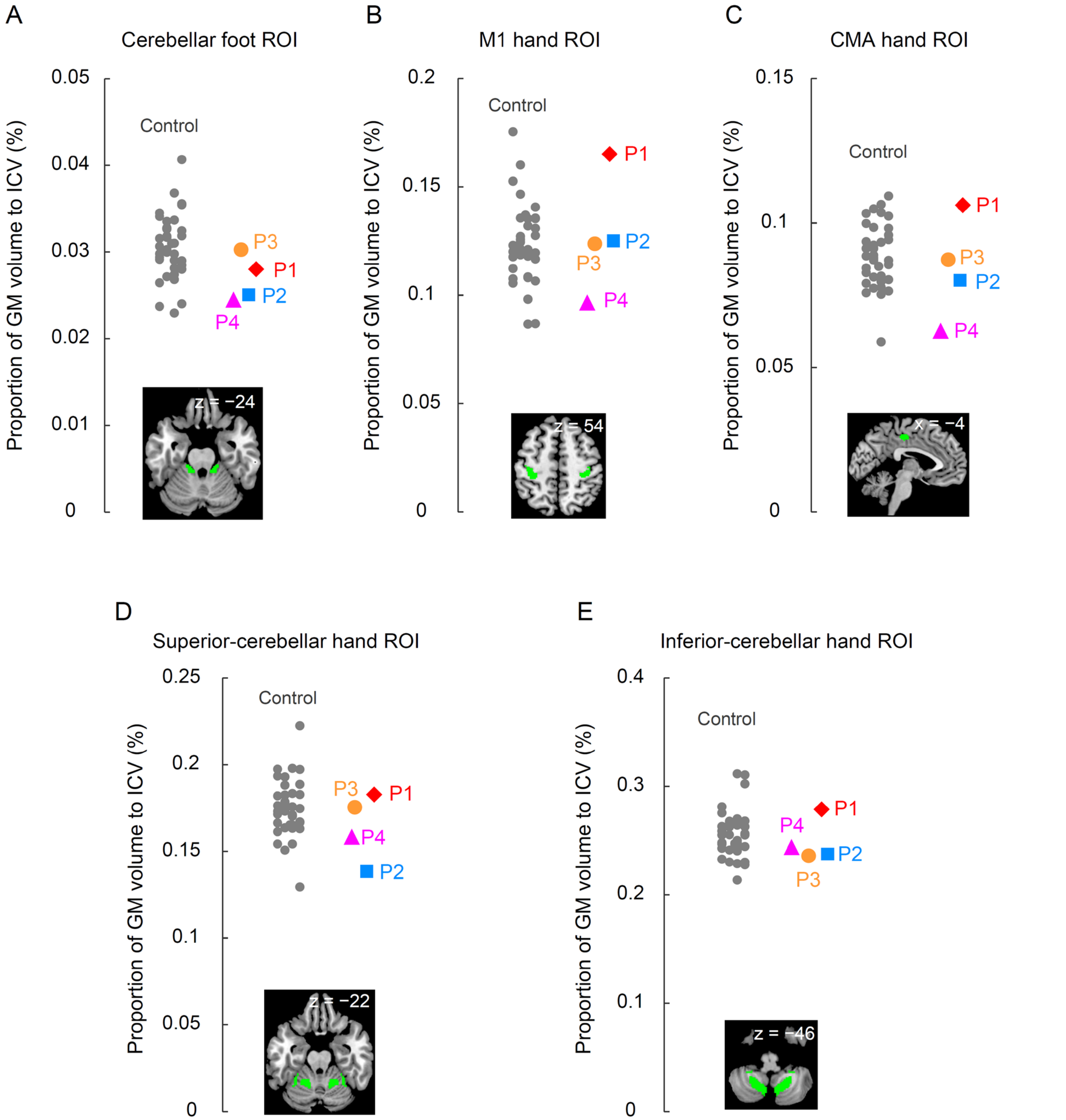


Figure S4. Individual GM volume in each of cerebellar foot ROI (A), the M1 hand ROI (B), CMA hand ROI (C), superior-cerebellar hand ROI (D), and inferior-cerebellar hand ROI (E). Each ROI is displayed at the bottom of each panel. In each panel, a vertical axis indicates the individual proportion of GM volume in each ROI to the ICV. Abbreviations: M1, primary motor cortex; CMA, cingulate motor area; ROI, region-of-interest; a.u., *arbitrary unit*; ICV, intracranial volume.
